# CDK1-dependent N-terminal NuMA phosphorylation promotes dynein-dynactin-NuMA assembly for accurate chromosome segregation

**DOI:** 10.1101/2025.07.19.665708

**Authors:** Marvin van Toorn, Keishi Shintomi, Tomomi Kiyomitsu

## Abstract

The microtubule-based motor dynein and its cofactor dynactin fulfil essential functions throughout the cell cycle, including organelle transport and mitotic spindle assembly. To achieve these diverse functions, dynein-dynactin associates with different activating adaptors. Nuclear Mitotic Apparatus (NuMA) is a mitosis-specific adaptor that connects dynein-dynactin with microtubules to focus mitotic spindle poles. NuMA’s C-terminal microtubule-binding activity is promoted by mitotic phosphorylation, but regulation of NuMA’s N-terminal interaction with dynein-dynactin remains unclear. Here, we combine a membrane-tethering assay, quantitative proteomics, and live functional analyses in human cells to show that the interaction between NuMA’s N-terminus and dynein-dynactin is cell cycle-regulated and driven by mitotic phosphorylation. We identify highly conserved CDK1 consensus sites proximal to NuMA’s dynein heavy chain-binding site, which are phosphorylated by CDK1-Cyclin B1 in cells and *in vitro*. This CDK1-dependent phosphorylation, together with NuMA’s Spindly-like motif, is crucial for stable dynein-dynactin-NuMA (DDN) complex formation. Replacement of endogenous NuMA with phosphorylation-deficient NuMA mutants leads to aberrant dynein distribution on mitotic spindles, resulting in chromosome mis-segregation and micronucleus formation. Together, our results highlight CDK1-dependent N-terminal NuMA phosphorylation as a crucial mitotic switch that constitutes a regulatable core of multivalent interactions with dynein-dynactin to assemble stable DDN complexes for accurate chromosome segregation.

## Introduction

During mitosis, a microtubule-based bipolar spindle is assembled to orchestrate segregation of duplicated chromosomes to daughter cells^1,2^. Formation and architecture of the spindle are controlled by spindle assembly factors (SAFs), which interact with microtubules and microtubule-associated proteins to spatially regulate microtubule nucleation, dynamics, transport, and crosslinking^3^. Defects in spindle assembly are associated with aneuploidy, cancer and birth defects, underscoring the importance of SAFs for human physiology^4^.

Nuclear Mitotic Apparatus (NuMA) is an abundant and well-established SAF that is highly conserved among vertebrates^5,6^. During interphase, NuMA localizes to the nucleus, where it contributes to nuclear formation, transcriptional regulation, DNA repair, and apoptosis^7,8^. In mitosis, NuMA regulates spindle assembly and bipolarization during prometaphase^9,10^. It also maintains microtubule minus-end focusing^11,12^, governs spindle orientation and positioning^13,14^ and tethers centrosomes to spindle poles^11,15^ during metaphase. NuMA deficiency during mitosis results in unfocused spindles, delayed and aberrant anaphase entry, and genomic instability through micronucleus formation^11,16,17^. After years of speculation, NuMA was recently confirmed to act as a *bona fide* activating adaptor for the minus-end-directed microtubule motor, cytoplasmic dynein-1 (hereafter dynein) *in vitro*^18,19^. Human dynein is a large complex comprised of pairs of catalytic heavy chains (DHC), intermediate chains (ICs), light intermediate (LICs) and light chains (LCs). This complex intrinsically adopts an autoinhibited (‘Phi’) conformation^20,21^. Indeed, dynein is dependent on its obligate co-factor dynactin, positive regulators such as LIS1, and NudE, and a collection of non-redundant activating adaptors to form fully functional dynein-dynactin-adaptor (DDA) complexes, thereby acquiring the processive motility to fulfill its numerous essential cellular functions^22–24^. These activating adaptors, which also include BICD2, Hook3, and Spindly, connect dynein to specific cargos, such as RNAs, protein complexes, or even entire organelles, to enable their microtubule-based transport^25^. While the adaptors share little overall sequence similarity, they do share certain structural elements, as they all form parallel dimers that contain a long central coiled-coil and one or multiple conserved dynein adaptor domains, namely Heavy chain binding sites (HBS1), CC1-boxes, Hook domains, and/or Spindly motifs. NuMA contains a Spindly-like motif (SpM)^13^ that associates with the pointed-end of dynactin^18^, as well as a CC1-like box and a Hook domain that interact with dynein LICs *in vitro*^26^. Additionally, an HBS1 site was recently identified in NuMA’s N-terminus^18^.

A central question in dynein biology is how dynein fulfills its numerous cellular roles in a dynamic and efficient manner, which requires its association with its various adaptors to be under tight spatiotemporal regulation. Most adaptors, including BICD1^27^, BICD2^28^, Hook3^29^ and Spindly^30^, adopt autoinhibited states that are relieved by cargo binding, suggesting that cargo availability may be a major regulator of dynein-adaptor interactions. However, other adaptors, including BICDL1^31^ and Rab11-FIP3^32^, bind dynein independently of their cargos, and whether NuMA can do so as well remains debated^18,19^. Interestingly, recent work indicates cell cycle-regulated BICD2 phosphorylation as an additional regulatory mechanism of dynein-dynactin-adaptor complex formation^33^. Mitotic NuMA undergoes extensive phosphorylation^34^ by various kinases, including CDK1^35,36^, PLK1^34,37^, Aurora A^14,38,39^ and ABL1^40^, affecting its localization, microtubule binding, and ability to position and orient the spindle. Intriguingly, however, most NuMA phosphorylation is reported in its unstructured C-terminal domain, whereas interaction with dynein-dynactin is mediated through the N-terminus. In fact, the first 705 amino acids of NuMA are sufficient for mitotic interaction with dynein^41^, as well as for cortical dynein recruitment during metaphase^13^. Thus, the exact link between NuMA phosphorylation and dynein-dynactin interaction remains contentious. Additionally, given that NuMA and dynein-dynactin can each associate independently with microtubules^7,12,42^, the precise contribution of their interaction to faithful chromosome segregation and genome stability remains to be fully determined.

Here, we show that the N-terminal domain of NuMA specifically interacts with dynein-dynactin during mitosis in human cells. Stable interaction with dynein-dynactin requires cell cycle-regulated phosphorylation of highly conserved serine residues located near NuMA’s HBS1 site, mediated by the mitotic CDK1 kinase. Failure to phosphorylate the N-terminus of NuMA during mitosis weakens the interaction with dynein-dynactin and leads to aberrant dynein distribution along the mitotic spindle, causing chromosome mis-segregation and micronucleus formation. Our results therefore highlight N-terminal NuMA phosphorylation as a crucial event to ensure genome stability during mitotic cell division.

## Results

### Cell cycle-dependent association of NuMA’s N-terminal region with dynein-dynactin

Since NuMA and dynein-dynactin are spatially separated by the nuclear membrane in the nucleus and cytoplasm, respectively^6,43,44^, we first determined whether formation of dynein-dynactin-NuMA (DDN) complexes is solely obstructed by this physical segregation. We previously reported that optogenetic targeting of NuMA-N(1-705) is sufficient for cortical dynein (DHC-SNAP) recruitment during metaphase^13^, but we found that this NuMA fragment was unable to recruit dynein to the membrane during interphase, in contrast to Hook3-N(1-552) **(Fig. S1a)**. This suggests that spatial separation of dynein-dynactin and NuMA is not the primary obstacle to DDN complex formation. To further explore the potential cell cycle-dependent association of NuMA with dynein-dynactin, we developed a membrane-tethering assay using Flp-In T-REx 293 cells^45^ **(Fig. 1a)**. N-terminal fragments of dynein adaptor constructs fused with mCherry-3xFlag and a plasma membrane-targeting CAAX motif (mCherry-3xFlag-CAAX) were conditionally expressed from the FRT site **(Fig. 1b)**, whereas endogenous dynein was visualized using mClover-tagged DHC (DHC-mClover, **Fig. S1b)**. We then examined the ability of these adaptors to recruit endogenous dynein during both interphase and mitosis **(Fig. 1a).** N-terminal fragments of the established dynein adaptors BICD2-N(1-400) and Hook3-N(1-552) successfully recruited DHC-mClover to the plasma membrane in interphase. However, a Hook3-N mutant (I154A) deficient in its ability to associate with dynein LIC1^29^ failed to recruit dynein **(Fig. 1c-d)**, showing the specificity of our experimental system. In stark contrast to Hook3-N and BICD2-N, NuMA-N(1-705) and Spindly-N(1-359) fragments failed to recruit dynein during interphase **(Fig. 1c-d)**, but selectively recruited dynein during mitosis **(Fig. 1e-f).** The interaction between NuMA-N and dynein appeared particularly potent, as it not only resulted in the greatest enrichment of cortical dynein, but also induced severe chromosome misalignments and robust prometaphase arrest, which were not observed with any other adaptor constructs **(Fig. 1e-g)**. This arrest phenotype was not due to variation in expression, as the CAAX-tagged adaptor constructs were expressed at similar levels **(Fig. S1c)**, and even cells with lower NuMA-N expression displayed mitotic arrest **(Fig. S1d)**. Moreover, NuMA-N also recruited the highest levels of dynein to the cortex in cells mitotically arrested with the Eg5-inhibitor, S-trityl-L-cysteine (STLC) **(Fig. S1e-f)**, suggesting that NuMA-N recruits dynein more efficiently, regardless of mitotic arrest status. We further confirmed the mitosis-specific interaction between NuMA-N and dynein-dynactin by immunoprecipitation. CAAX-tagged NuMA-N preferentially interacted with DHC and the dynactin subunit, DCTN1 (p150^Glued^), in STLC-treated prometaphase cells **(Fig. 1h)**, whereas BICD2-N associated with these proteins in both interphase (arrested at G2/M boundary with the CDK1 inhibitor RO3306) and in mitotic cell extracts. Together, these data show that DDN complex formation is not solely hindered by their physical separation between the nucleus and cytoplasm, but is also subject to cell cycle-dependent regulation.

**Figure 1:**
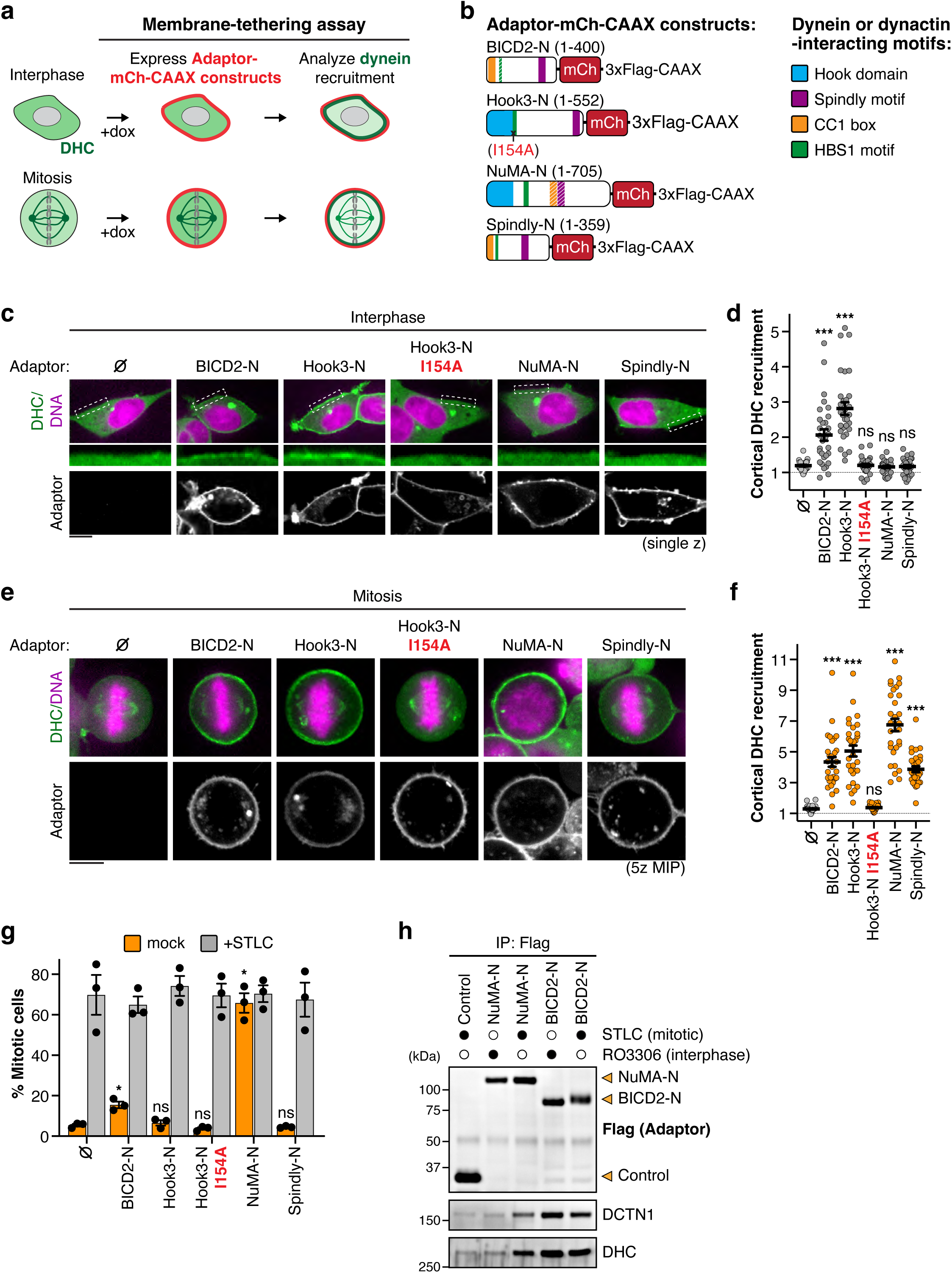
Cell cycle-dependent association between NuMA-N and dynein-dynactin. **a)** Schematic illustrating the principle of the doxycycline (Dox)-inducible N-terminal dynein adaptor tethering assay in DHC-mClover knock-in (KI) Flp-In T-REx 293 cells and the resulting dynein-dynactin-adaptor (DDA) complex assembly at the membrane. **b)** Diagram showing adaptor constructs used and their identified dynein-dynactin interacting domains. **c)** Representative live cell images showing endogenous dynein (DHC-mClover) recruitment to the indicated membrane-tethered adaptor constructs in the presence of 2 μg/mL Dox during interphase. **d)** Quantification of DHC-mClover accumulation at the membrane relative to its cytoplasmic signal. n = 30 cells ± SEM from three independent experiments; Brown-Forsythe and Welch ANOVA with Dunnett’s multiple comparisons tests. **e)** Representative images of cortical dynein recruitment by the indicated adaptor constructs during mitosis. **f)** Quantification of DHC-mClover accumulation at the membrane relative to its cytoplasmic signal. n = 30 cells ± SEM from three independent experiments; Brown-Forsythe and Welch ANOVA with Dunnett’s multiple comparisons tests. **g)** Quantification of the percentage of mitotic cells after expression of the indicated N-terminal adaptor constructs. 10 μM STLC was added to cells together with 2 μg/mL Dox for overnight incubation where indicated. n = 3 experiments ± SEM; Brown-Forsythe and Welch ANOVA with Dunnett’s multiple comparisons tests. **h)** Flag immunoprecipitation of mCherry-3xFlag-CAAX alone (Control) or conjugated to BICD2-N and NuMA-N constructs in either interphase-(RO3306-treated) or mitosis-arrested (STLC-treated) DHC-mClover KI Flp-In T-REx 293 cells followed by immunoblotting for the indicated proteins. Scale bar represents 10 μm; MIP: maximum intensity projection.

### Mitotic interactions and phosphorylation of NuMA and dynein-dynactin

To analyze whether mitotic modifications govern endogenous DDN complex formation and dissolution, we next optimized conditions to purify the DDN complex from either asynchronously growing or mitotically-arrested (STLC-treated) cell extracts using hypotonic or dithiobis(succinimidyl propionate) [DSP]-crosslinked immunoprecipitation. Hypotonic immunoprecipitation of endogenous DCTN2/p50-mAID-mClover-3xFlag (DCTN2-mACF, co-expressed with DCTN1/p150-SNAP, **Fig. S2a-b**) confirmed constant interactions with DCTN1 and DHC throughout the cell cycle **(Fig. 2a),** whereas full-length Hook3 preferentially interacted with DCTN2 during interphase. Under these conditions, NuMA was only solubilized in mitotic cells due to the inability of digitonin to disrupt nuclear membranes, but efficiently co-immunoprecipitated with DCTN2 during mitosis. Similar results were obtained when DCTN2-mACF was immunoprecipitated under crosslinked conditions, in which NuMA was lysed in both interphase and mitosis, but co-immunoprecipitated with DCTN2 only in mitosis^46^ **(Fig. 2b)**. Additionally, crosslinked NuMA-mACF immunoprecipitation showed mitotic interactions with DHC, DCTN1 and its cortical docking protein LGN **(Fig. 2c),** further validating our approaches for detecting cell cycle-dependent interactions of NuMA and dynein-dynactin.

**Figure 2:**
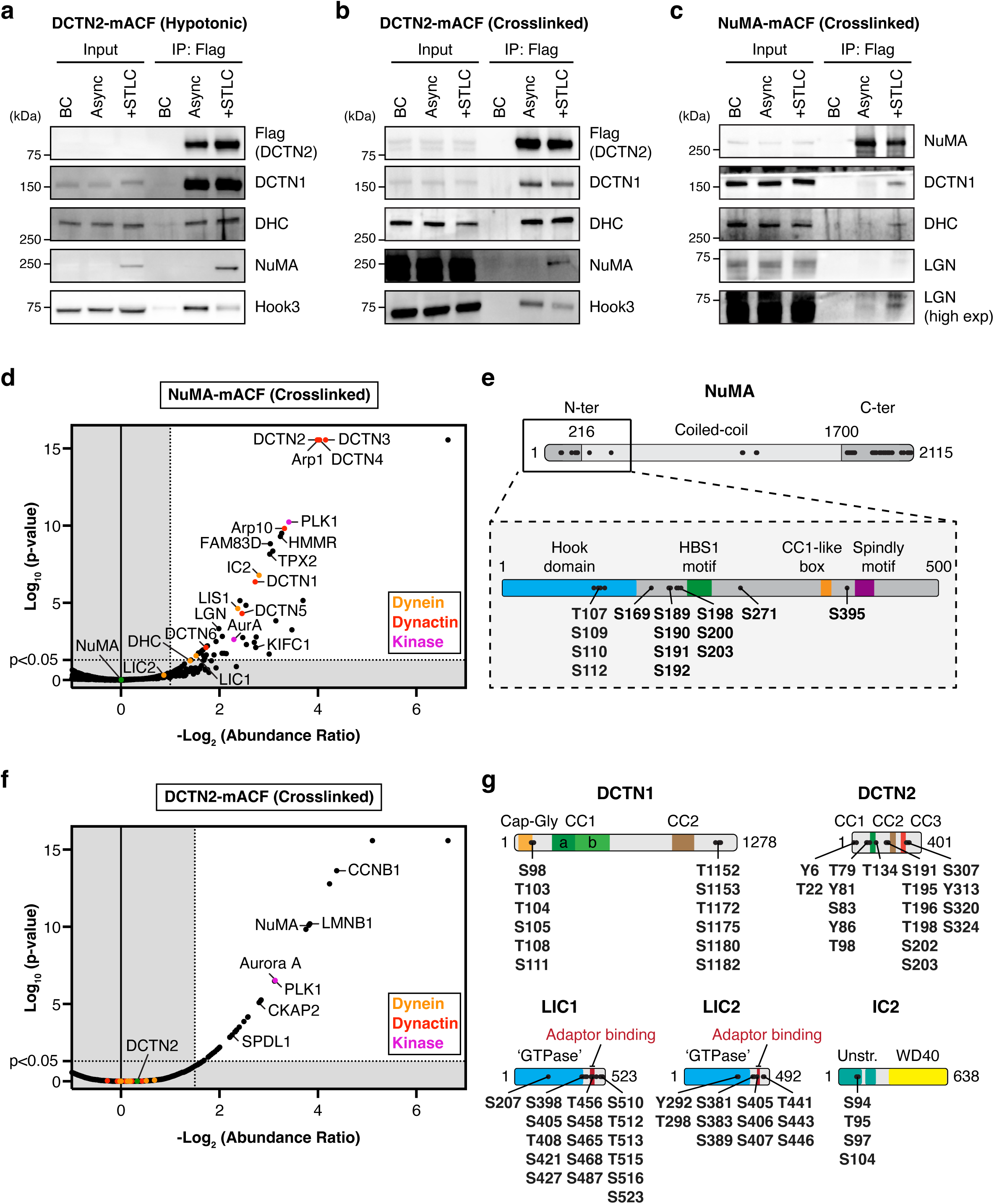
Identification of mitosis-specific DDN interactors and modifications. **a)** Hypotonic Flag immunoprecipitation of DCTN2-mACF from either asynchronously growing or mitosis-arrested (STLC-treated) HCT116 DCTN2-mACF / DCTN1-SNAP KI cell lysates, followed by immunoblotting for indicated proteins. BC: Binding control. **b)** Crosslinked Flag immunoprecipitation of DCTN2-mACF from asynchronous or mitosis-arrested (STLC-treated) HCT116 DCTN2-mACF / DCTN1-SNAP KI cell lysates, followed by immunoblotting for the proteins shown. **c)** Crosslinked Flag immunoprecipitation NuMA-mACF in either asynchronously growing or mitosis-arrested (STLC-treated) HCT116 NuMA-mACF KI cells, followed by immunoblotting for the indicated proteins. **d)** Volcano plot highlighting mitosis-specific NuMA binding partners after crosslinked IP from either asynchronously growing or prometaphase-arrested (STLC-treated) HCT116 NuMA-mACF KI cells. Eluates were digested and analyzed by mass spectrometry followed by label-free quantification using the CHIMERYS search algorithm. **e)** Schematic overview of NuMA phosphorylation sites identified using crosslinked immunoprecipitation followed by mass spectrometry and the SequestHT search algorithm. **f)** Volcano plot showing mitosis-specific DCTN2 binding partners identified by crosslinked IP from either asynchronously growing or prometaphase-arrested (STLC-treated) HCT116 DCTN2-mACF KI cells. Eluates were digested and analyzed by mass spectrometry followed by label-free quantification using the CHIMERYS search algorithm. **g)** Schematic overview of dynactin (DCTN1 and DCTN2) and dynein (LIC1, LIC2 and IC2) subunit phosphorylation sites revealed by crosslinked immunoprecipitation-mass spectrometry and subsequent analysis through the SequestHT search algorithm.

We then expanded these results using label-free quantitative proteomics to more comprehensively compare cell cycle-regulated interactions and modifications of NuMA, dynein, and dynactin **(Fig. S2c).** We employed crosslinked immunoprecipitation followed by mass spectrometry to obtain the most complete and unbiased overview of changes in the interactomes. Crosslinked NuMA immunoprecipitation validated its mitotic interaction with the majority of dynactin and dynein subunits, the dynein regulator LIS1, and other previously identified binding partners such as ASPM^47^, KIFC1^10^ and LGN^48^ **(Fig. 2d, Table S1).** In addition, mitotic NuMA associated with the well-established mitotic kinases Aurora A and PLK1, as well as the spindle-positioning proteins, HMMR and FAM83D, which form a complex with CK1α kinase^49^. These mitotic interactions were corroborated using an alternative biotin-based proximity labeling^50^ approach with endogenous NuMA-mClover-miniTurbo, which additionally revealed proximal localization of CK1α, CDK1 and its activating partner Cyclin B1 **(Fig. S2d-g, Table S2)**. Importantly, we also found multiple mitosis-specific phosphorylation sites using crosslinked immunoprecipitation. While the majority of phosphorylation sites were located at the C-terminus of NuMA, several were located on its N-terminal domain **(Fig. 2e, Table S1)**, particularly upstream of coiled-coil CC1a (aa 212-269) and the recently identified dynein heavy chain-binding site 1 (HBS1, aa 212-239)^18^ embedded herein.

Mass spectrometry analysis of DCTN2 interactions under crosslinked conditions revealed consistent interaction with most core dynein-dynactin subunits, including LIS1, during interphase and mitosis **(Fig. 2f, Table S3).** Mitotic dynein-dynactin specifically interacted with Lamin B1^51^, its mitotic adaptors NuMA and Spindly, the mitotic kinases Aurora A and PLK1 and the CDK1-regulator Cyclin B1 **(Fig. 2f)**. Like NuMA, dynein and dynactin underwent cell cycle-regulated phosphorylation on DCTN1, DCTN2, DHC, LIC1 and LIC2 subunits **(Fig. 2g, Table S3).** Collectively, these data reveal cell cycle-dependent interaction partners and phosphorylation of NuMA and dynein-dynactin, suggesting that cell cycle-dependent modifications are involved in DDN complex formation.

### CDK1 kinase activity stabilizes DDN assembly in mitosis

Intrigued by the cell cycle-dependent phosphorylation of NuMA and dynein-dynactin, we next investigated whether inhibition of known (Aurora A, PLK1 and CDK1) or newly identified (CK1α) kinases, detected by our proteomic studies, affected association of NuMA-N with dynein-dynactin during mitosis using our membrane-targeting assay. Importantly, live cell imaging showed that inhibition of either CDK1 or PLK1 using the chemical inhibitors, RO3306^52^, and BI2536^53^, respectively, markedly decreased cortical dynein recruitment by membrane-associated NuMA-N during mitosis **(Fig. 3a-b).** Conversely, Aurora A inhibition by MLN8237^54^ did not reduce cortical dynein recruitment, whereas D4476, a selective CK1 inhibitor^55^, only slightly reduced cortical dynein accumulation despite extended incubation **(Fig. S3a-b)**.

**Figure 3:**
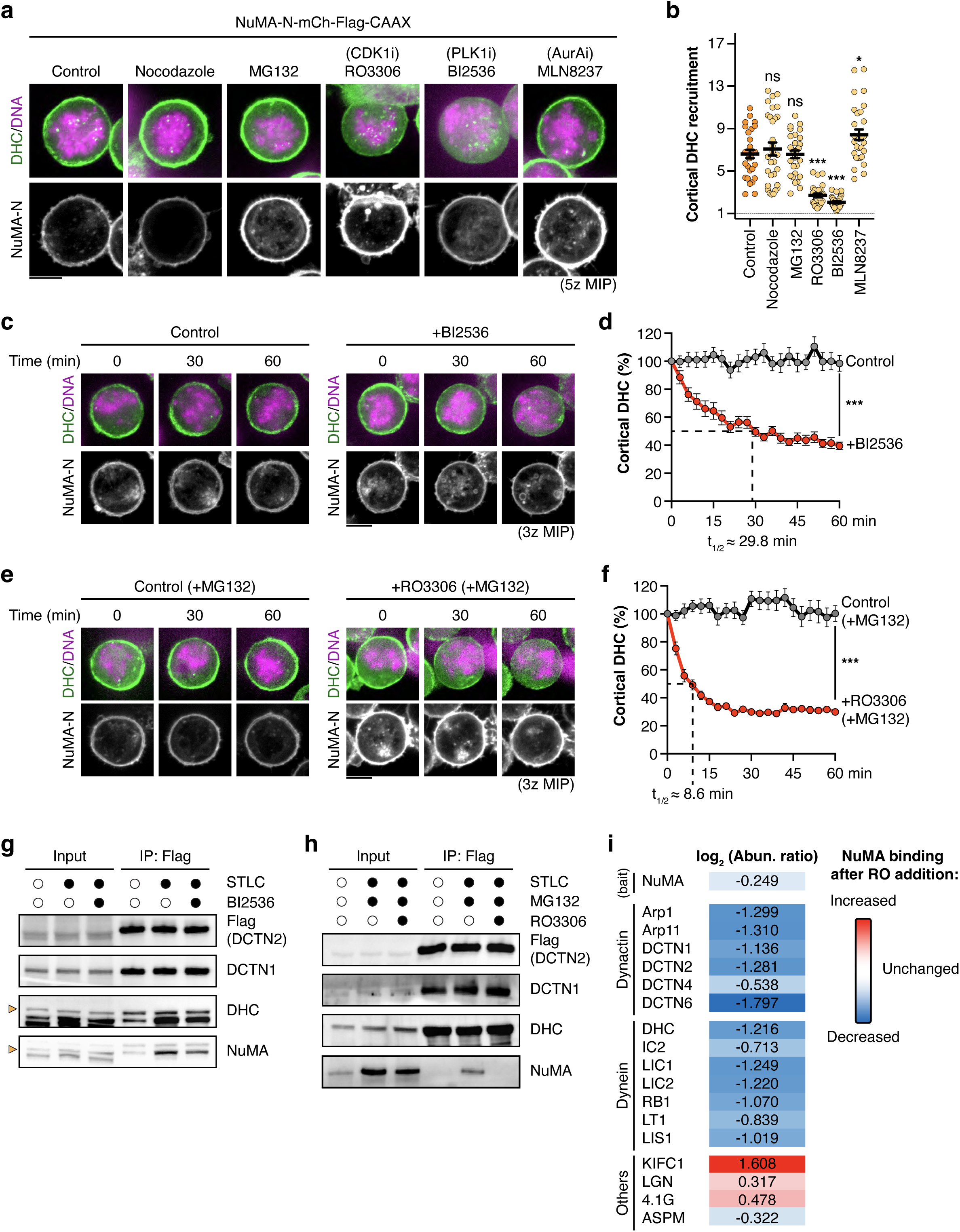
CDK1 activity drives DDN complex assembly. **a)** Representative live cell images showing endogenous dynein (DHC-mClover) recruitment to membrane-tethered NuMA-N in Flp-In T-REx 293 cells after treatment with the selected inhibitors. Inhibitors were administered for 2 h, except for RO3306 which was added for <30 min to avoid mitotic exit. **b)** Quantification of relative DHC-mClover accumulation at the membrane. n = 30 cells ± SEM from three independent experiments; Brown-Forsythe and Welch ANOVA with Dunnett’s multiple comparisons tests. **c)** Representative images showing time-lapse imaging of NuMA-N-dependent cortical DHC-mClover accumulation in Flp-In T-REx 293 cell after addition of 10 μM BI2536 or 0.1% DMSO as a control. **d)** Quantification of cortical dynein at indicated time points, normalized to 100% at t = 0 min. n = 24 cells ± SEM from four independent experiments; unpaired Welch’s t-test on the final values (t = 60 min). **e)** Representative images showing time-lapse imaging of NuMA-N-dependent cortical DHC-mClover accumulation in Flp-In T-REx 293 cell after addition of 9 μM RO3306 (1 h after pre-incubation with 20 μM MG132) or MG132 and 0.1% DMSO as a control. **f)** Quantification of cortical dynein at indicated time points, normalized to 100% at t = 0 min. n = 24 cells ± SEM from four independent experiments; unpaired Welch’s t-test on the final values (t = 60 min). **g-h)** Hypotonic Flag immunoprecipitation of DCTN2-mACF in either asynchronously growing or mitosis-arrested (STLC-treated) HCT116 DCTN2-mACF / DHC-SNAP KI cells followed by immunoblotting for the proteins shown. Where indicated, 10 μM BI2536 (1 h) or 20 μM MG132 (1 h pre-incubation) and 9 μM RO3306 (15 min) were added to cells prior to collection and lysis. **i)** Heatmap showing log_2_-transformed abundance ratios of mitotic NuMA interactors after incubation with RO3306 for 15 min (in the presence of 20 μM MG132) relative to control (mitotic NuMA interactors in the presence of MG132 alone). A positive value represents increase in interaction after RO3306, whereas a negative value represents a decrease. Ratios were calculated from three independent experiments. Scale bars represent 10 μm.

Since these data suggested that CDK1 and/or PLK1 are the major kinases governing dynein-dynactin-NuMA complex formation, we subsequently quantified dissociation kinetics of cortical dynein after inhibition of these kinases in real time. We observed that PLK1 inhibition gradually reduced cortical dynein, reaching half intensity (t_1/2_) after ∼ 29.8 min **(Fig. 3c-d).** Similar effects also occurred with nanomolar concentrations of either BI2536 (t_1/2_ ≈ 20.7 min) or GSK46136 (t_1/2_ ≈ 29.5 min), the latter of which specifically inhibits PLK1 **(Fig. S3c-d).** Notably, CDK1 inhibition by RO3306 triggered a more rapid (t_1/2_ ≈ 13.7 min) and robust dynein dissociation from membrane-bound NuMA-N **(Fig. S3e)**, even after pre-incubation with the proteasome inhibitor, MG132 (t_1/2_ ≈ 8.6 min, **Fig. 3e-f)** to delay mitotic slippage by preventing Cyclin B degradation^56^. In contrast, short-term CDK1 inhibition did not significantly impair cortical dynein recruitment by Hook3-N or Spindly-N **(Fig. S3f-g),** indicating that CDK1 specifically affects DDN complex formation.

We next examined whether these kinases also affected mitotic association of full-length NuMA with dynein-dynactin. We performed hypotonic immunoprecipitation of endogenous DCTN2-mACF and found that PLK1 inhibition modestly reduced the interaction between dynein-dynactin and NuMA, as residual interaction was still detectable even after 1 hr of PLK1 inhibition **(Fig. 3g).** In contrast, CDK1 inhibition in the presence of MG132 fully dissociated endogenous NuMA from dynein-dynactin within 15 min **(Fig. 3h).** Reciprocal immunoprecipitation of NuMA-mACF in the presence of RO3306 and MG132 followed by quantitative mass spectrometry also revealed dissociation of multiple dynein-dynactin subunits upon RO3306 treatment, whereas other mitotic NuMA interactors such as LGN, 4.1G, ASPM and KIFC1 remained associated with NuMA **(Fig. 3i).** In summary, our results indicate that CDK1 specifically stabilizes DDN complex assembly during mitosis.

### Phosphorylation of CDK1 consensus sites on NuMA-N promotes DDN assembly

Our mass spectrometry data identified 13 high-confidence phosphorylation sites in the first 300 amino acids of NuMA-N **(Fig. 2e).** To identify phosphorylation sites regulating DDN assembly, we mutated several of these serine/threonine residues to alanine and evaluated their effects on cortical dynein accumulation and the associated mitotic arrest phenotype using our membrane-tethering assay. Intriguingly, S198A, S200A and S203A (henceforth referred to as S3A) mutations, but not other mutations, significantly alleviated the chromosome misalignment and mitotic arrest induced by NuMA-N **(Fig. 4a-b)**. Although cells expressing NuMA-N S3A displayed similar cortical dynein intensities **(Fig 4c)** these cells occasionally entered anaphase with aberrations such as chromosome bridges or lagging chromosomes **(Fig. S4a)**, leading to micronucleus formation during interphase **(Fig. S4b).**

**Figure 4:**
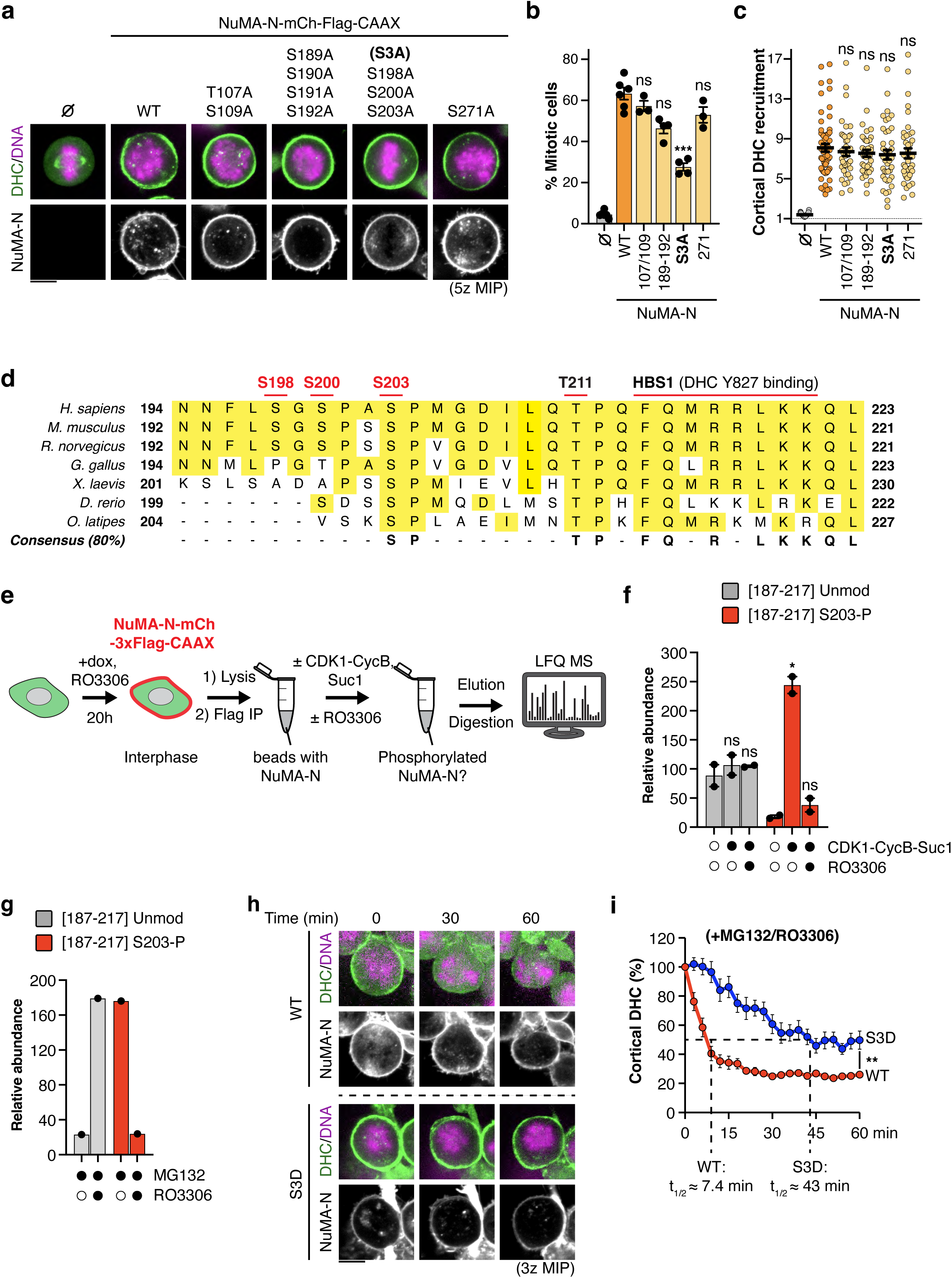
CDK1-mediated NuMA phosphorylation adjacent to the HBS1 site regulates dynein binding. **a)** Representative images showing endogenous dynein (DHC-mClover) recruitment to the indicated membrane-tethered NuMA-N phosphomutants. **b)** Quantification of the percentage of mitotic cells after expression of the displayed N-terminal NuMA constructs. **c)** Quantification of relative DHC-mClover accumulation at the membrane in mitotic cells. n = 30 cells ± SEM from three independent experiments; Brown-Forsythe and Welch ANOVA with Dunnett’s multiple comparisons tests. **d)** Sequence alignment showing conservation of phosphorylated residues adjacent to the heavy chain binding site (HBS1) within the N-terminus of NuMA. **e)** Schematic showing the workflow of CDK1-mediated NuMA-N phosphorylation *in vitro* and detection. **f)** Mass spectrometry-based quantification either unmodified or S203-phosphorylated NuMA [187-217 aa] peptides after incubation with CDK1-CycB1-Suc1 and RO3306. n = 2 independent experiments ± SEM **g)** Mass spectrometry-based quantification either unmodified or S203-phosphorylated NuMA [187-217 aa] peptides isolated from mitotic NuMA-N-mCherry-3xFlag-CAAX expressing Flp-In T-REx 293 cells. Where indicated, 9 μM RO3306 was added in addition to 20 μM MG132 for 15 min. n = 1 experiment. **h)** Top: Representative images of time-lapse imaging of wild-type or phosphomimetic (S3D) NuMA-N-dependent cortical DHC-mClover accumulation in Flp-In T-REx 293 cells after addition of 9 μM RO3306 (1 h after pre-incubation with 20 μM MG132). **i)** Quantification of cortical dynein at indicated time points, normalized to 100% at t = 0 min. n = 18 cells ± SEM from three independent experiments; unpaired Welch’s t-test on the final values (t = 60 min). Scale bars represent 10 μm.

The phosphorylation sites (S198, S200, and S203) mutated in NuMA-N S3A were highly conserved (**Fig. 4d**) and localized upstream of the DHC-binding HBS1 domain^18^, suggesting that their phosphorylation may stimulate interaction between HBS1 and DHC. Because NuMA-N S3A only partially alleviated the mitotic arrest phenotype (**Fig. 4b**), we tested whether additional mutagenesis of the similarly evolutionary conserved threonine 211 (T211), located within 5 amino acids of the HBS1 domain, had similar or additional effects. However, mutagenesis of T211 did not further alleviate the mitotic arrest phenotype of NuMA-N S3A, nor did it have any effect on its own (Fig. S4c-e), indicating that T211 is not involved in DDN assembly.

S200 and S203 in NuMA-N are both consistent with the CDK1 consensus (S/T-P)^57,58^. To test whether these residues are indeed phosphorylated by CDK1, we purified NuMA-N from interphase cells, incubated it with recombinant CDK1-Cyclin B1-Suc1, capable of multi-site phosphorylation *in vitro*^59^, and determined the phosphorylation status of NuMA-N by mass spectrometry (**Fig. 4e**). We found that S203 of NuMA-N was phosphorylated upon incubation with CDK1-Cyclin B1-Suc1, but not when the CDK1 inhibitor RO3306 was added to the reaction mixtures as well (**Fig. 4f**). Moreover, incubation with RO3306 also decreased S203 phosphorylation in mitotic cells expressing NuMA-N (**Fig. 4g**). Together, these data confirm that CDK1 is responsible for mitotic NuMA phosphorylation of at least S203, both in cells and *in vitro*.

To establish whether N-terminal NuMA phosphorylation alone is sufficient to drive DDN complex assembly in cells, we generated a phosphomimetic NuMA-N mutant in which S198, S200 and S203 were replaced with aspartic acid residues (S3D). Although most cells expressing this NuMA-N S3D construct were negative for cortical dynein accumulation during interphase, some cells showed weak dynein signals at the membrane (Fig. S4f). To determine whether this could be explained by a slower release from dynein-dynactin upon CDK1 inactivation during mitotic exit^60^, we performed time-lapse imaging upon addition of RO3306 (in the presence of MG132) and tracked the reduction in cortical dynein accumulation. NuMA-N S3D (t_1/2_ ≈ 43.0 min) showed a significantly reduced rate of dynein dissociation from the cortex upon RO3306 administration compared to NuMA-N WT (t_1/2_ ≈ 7.4 min) (**Fig. 4h-i**). Together, these findings suggest that although N-terminal phosphorylation alone is insufficient for DDN assembly, timely phosphorylation and dephosphorylation of N-terminal NuMA residues are important in triggering the assembly and disassembly of the DDN complex.

### N-terminal NuMA phosphorylation and the Spindly-like motif cooperatively promote interaction with dynein-dynactin

Besides the HBS1 domain, NuMA’s N-terminus uniquely contains three additional dynein-dynactin binding motifs: a Hook domain, a CC1-like box, and a Spindly-like motif^6^ **(Fig. 1b).** Since many adaptors bind dynein-dynactin through multivalent interactions^25^, we next tested which of these domains could stimulate dynein-dynactin binding in cooperation with N-terminal NuMA phosphorylation. Our membrane-targeting assay showed that mutation of the Spindly-like motif (SpM) strongly reduced mitotic arrest induced by NuMA-N expression, whereas mutation of the CC1-like box or the Hook domain did not **(Fig. 5a-c and S5a-c**), consistent with recent structural analysis of the DDN complex^18^. Notably, NuMA-N carrying mutations at both the phosphorylation and Spindly-like motif (S3A/SpM) sites showed an additional reduction in cortical dynein intensities **(Fig. 5c).** Concomitantly, cells expressing this NuMA-N S3A/SpM construct showed increased levels of spindle pole-associated dynein, compared to cells expressing NuMA-N S3A or NuMA-N SpM, indicating impaired sequestration to the cell cortex **(Fig. 5d-e).** In conclusion, these data reveal that NuMA’s dynactin-interacting Spindly-like motif cooperates with CDK1-mediated phosphorylation near the dynein-binding HBS1 site to stabilize the interaction between NuMA and dynein-dynactin.

**Figure 5:**
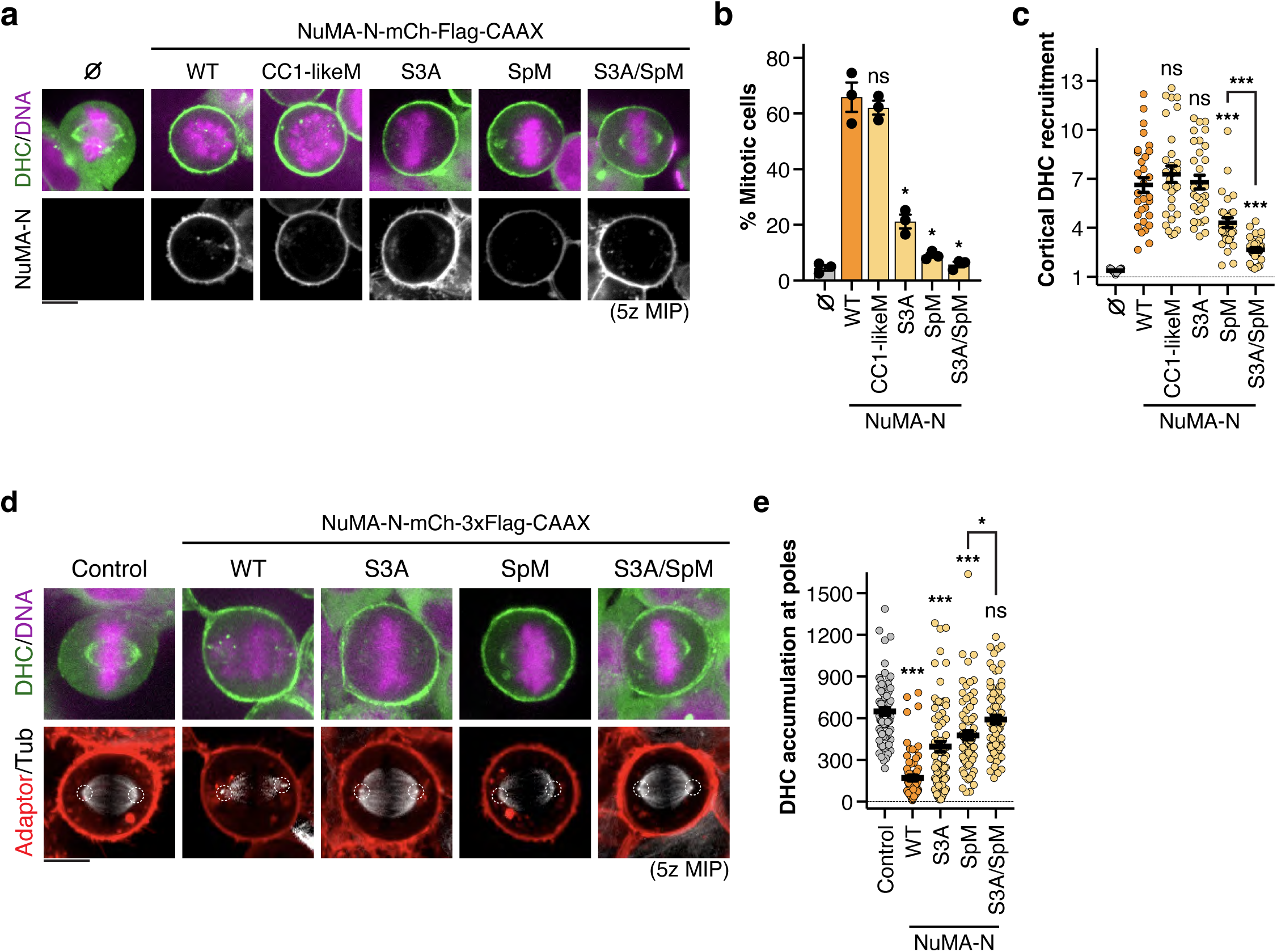
N-terminal NuMA phosphorylation and the Spindly-like motif cooperatively stimulate dynein binding. **a)** Representative images of cortical dynein recruitment by the indicated NuMA-N constructs during mitosis. **b)** Percentage of mitotic cells after expression of the displayed N-terminal NuMA constructs. **c)** Quantification of DHC-mClover accumulation at the membrane relative to its cytoplasmic signal. n = 30 cells ± SEM from three independent experiments; Brown-Forsythe and Welch ANOVA with Dunnett’s multiple comparisons tests. **d)** Representative images of dynein recruitment at spindle poles (visualized by SiR700-Tubulin) in mitotic Flp-In T-REx 293 DHC-mClover KI expressing the indicated NuMA-N mutants, or mCherry-3xFlag-CAAX alone (Control). **e)** Quantification of DHC-mClover accumulation at spindle poles. n = 72 spindle poles ± SEM from three independent experiments; Brown-Forsythe and Welch ANOVA with Dunnett’s multiple comparisons tests. Scale bars represent 10 μm.

### N-terminal phosphorylation sites and the Spindly-like motif are required for NuMA’s mitotic functions

We previously reported that NuMA depletion by auxin-inducible degron (AID) technology generates micronuclei due to mitotic chromosome mis-segregation in HCT116 cells^11^. To test whether N-terminal NuMA phosphorylation contributes to proper chromosome segregation, we first determined whether expression of phospho-deficient NuMA could rescue micronucleus formation caused by endogenous NuMA depletion using a replacement strategy **(Fig. S6a-b).** Expression of mCherry-NuMA WT after degradation of endogenous NuMA nearly completely prevented micronucleus formation 24 h after inducing endogenous NuMA degradation, whereas expression of either phospho-deficient (S3A) or Spindly-like motif mutant (SpM) NuMA only partially alleviated this phenotype. A S2A phospho-deficient mutant (S200A and S203A) showed a similar effect as when S198 was also mutated (S3A), consistent with S200 and S203 as primary phosphorylation sites. Importantly, in keeping with our results using NuMA-N **(Fig. 5)**, full length NuMA with doubly mutated (S3A/SpM) NuMA rescued micronucleus formation less effectively than either mutation alone, confirming that the N-terminal phosphorylation sites and the Spindly-like motif of NuMA cooperatively regulate functions of full-length NuMA **(Fig. 6a-b)**.

**Figure 6:**
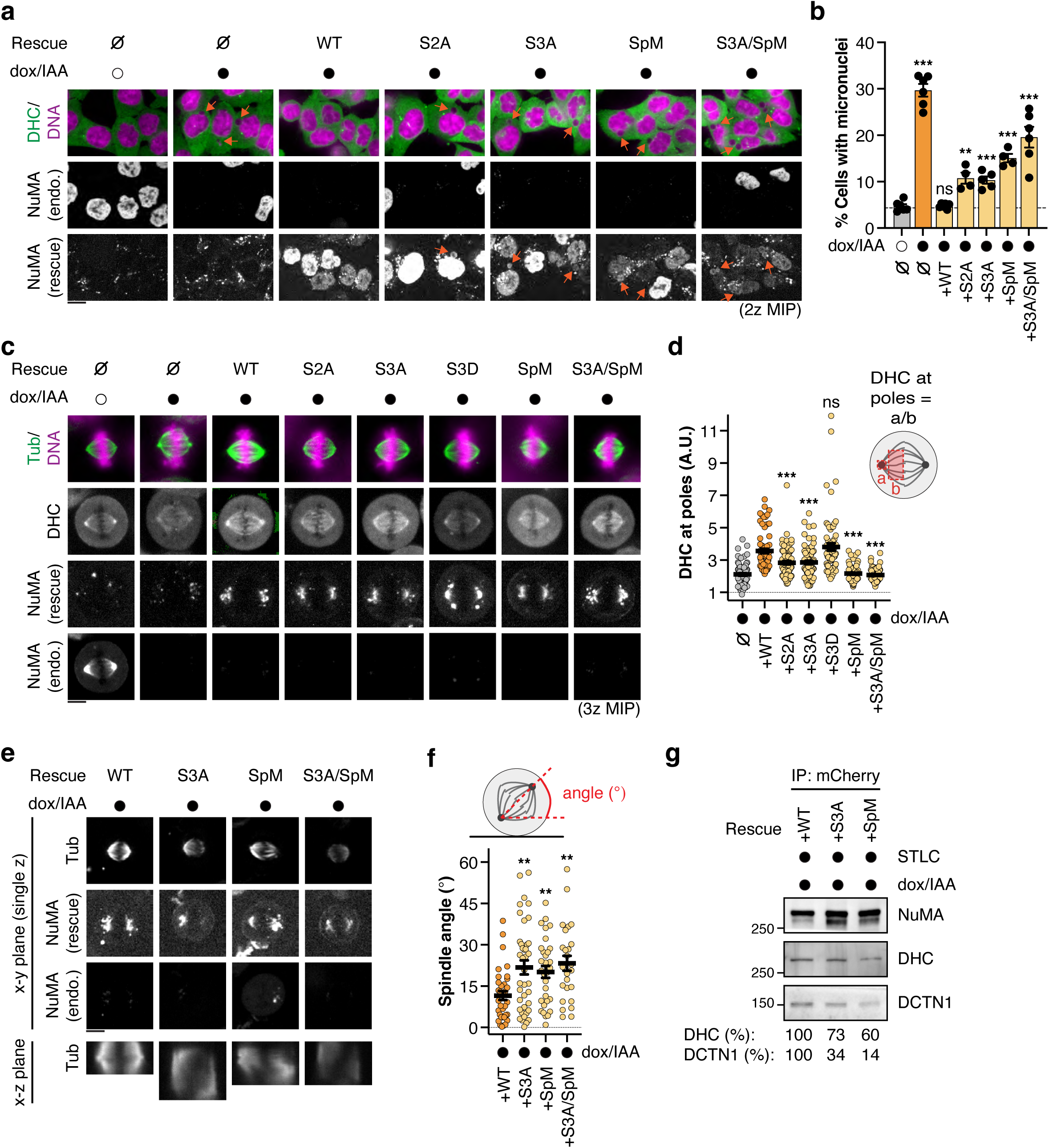
DDN functionality depends on N-terminal NuMA phosphorylation. **a)** Representative live cell fluorescent images showing the presence of micronuclei (orange arrowheads) in interphase NuMA-mACF / DHC-SNAP double-KI HCT116 cells. Where indicated, endogenous NuMA-mACF was depleted and replaced with the indicated mCherry-NuMA constructs through addition of Dox and IAA for 20 h. **b)** Quantification of micronucleus frequency. n > 4 independent experiments ± SEM; Brown-Forsythe and Welch ANOVA with Dunnett’s multiple comparisons tests. **c)** Representative live cell fluorescent images of endogenous dynein (DHC) distribution on the mitotic spindle in metaphase HCT116 NuMA-mACF / DHC-SNAP double-KI cells in which endogenous NuMA was replaced by the indicated NuMA mutants. **d)** Quantification of the ratio of pole dynein/spindle dynein intensities. n > 66 spindle poles from 3 independent experiments ± SEM; Brown-Forsythe and Welch ANOVA with Dunnett’s multiple comparisons tests. **e)** Orthogonal view of metaphase spindles in the x-y (top) and x-z (bottom) planes after NuMA replacement through Dox/IAA treatment for 24 h. **f)** Scatterplots of spindle angles on the x-z plane. n > 27 spindles from 2 independent experiments; Brown-Forsythe and Welch ANOVA with Dunnett’s multiple comparisons tests. **g)** Hypotonic immunoprecipitation of mitotic mCherry-NuMA (WT, S3A and SpM) from STLC-arrested prometaphase cells treated with Dox and IAA overnight, followed by immunoblotting for the indicated proteins. Relative DHC and DCTN1 binding (normalized to NuMA) are given below. Scale bars represent 10 μm.

Micronuclei can result from defects in mitotic spindle structure and/or function^61^. Therefore, we next analyzed metaphase spindles for abnormalities, particularly focusing on dynein localization. In control (-Dox/IAA) cells, punctate NuMA and dynein (visualized by endogenous DHC-SNAP^13^) signals were readily detected at spindle poles, likely representing DDN complexes actively transporting and clustering the minus-ends of spindle microtubules at the poles^12^ **(Fig. 5c).** As previously observed for dynactin^12^, dynein localization at spindle poles was markedly reduced upon NuMA depletion and instead exhibited a more diffuse distribution along the spindle **(Fig. 5c-d)**. Expression of WT or phosphomimetic (S3D) NuMA partially restored DHC accumulation at spindle poles, while NuMA S2A, S3A, SpM and S3A/SpM did not, even though these mutants accumulated at the poles like NuMA WT **(Fig. 5c-d).** Orientation of metaphase spindles, which is regulated through cortical DDN^13,41^, was also dysregulated after replacing endogenous NuMA with phospho-deficient or Spindly-like motif mutant NuMA **(Fig. 5e-f**). Furthermore, immunoprecipitation of either mCh-NuMA WT or mutants from mitotic cells using hypotonic conditions showed that NuMA S3A or SpM mutants exhibited reduced binding to both dynein and dynactin **(Fig. 5g)**. Together, these data demonstrate that N-terminal phosphorylation and Spindly-like motif enable stable association of full-length NuMA with dynein-dynactin in mitotic cells, thereby regulating dynein enrichment at spindle poles and spindle orientation.

### N-terminal NuMA phosphorylation and the Spindly-like motif are required for accurate chromosome segregation

To analyze the significance of N-terminal NuMA phosphorylation and the Spindly-like motif in chromosome segregation during anaphase, we performed STLC-arrest and release experiments while replacing endogenous NuMA-mACF with either mCherry-NuMA WT or mutants **(Fig. 7a).** When endogenous NuMA was replaced with NuMA WT, most cells showed successful metaphase chromosome alignment after release from STLC-induced prometaphase arrest and subsequently entered anaphase, on average, ∼18 min later **(Fig. 7b)**. Anaphase entry timing from metaphase was not significantly altered by replacement of endogenous NuMA-mACF with NuMA SpM **(Fig. 7b)**. However, the majority of cells exhibited chromosome bridges or lagging chromosomes during anaphase, phenotypes also associated with NuMA depletion^11,17^ and micronucleus formation^61^ **(Fig. 7c).** Like NuMA-SpM, NuMA-S3A replacement also increased the rate of chromosome mis-segregation after STLC washout, whereas anaphase entry timing was unaffected **(Fig. 7b, d)**, indicating that the spindle assembly checkpoint was intact and that these phenotypes were the result of spindle defects rather than prolonged mitosis. Together, these data indicate that N-terminal NuMA phosphorylation and the Spindly-like motif modulate dynein’s function in the spindle to enable accurate chromosome segregation and prevent micronucleus formation during mitosis.

**Figure 7:**
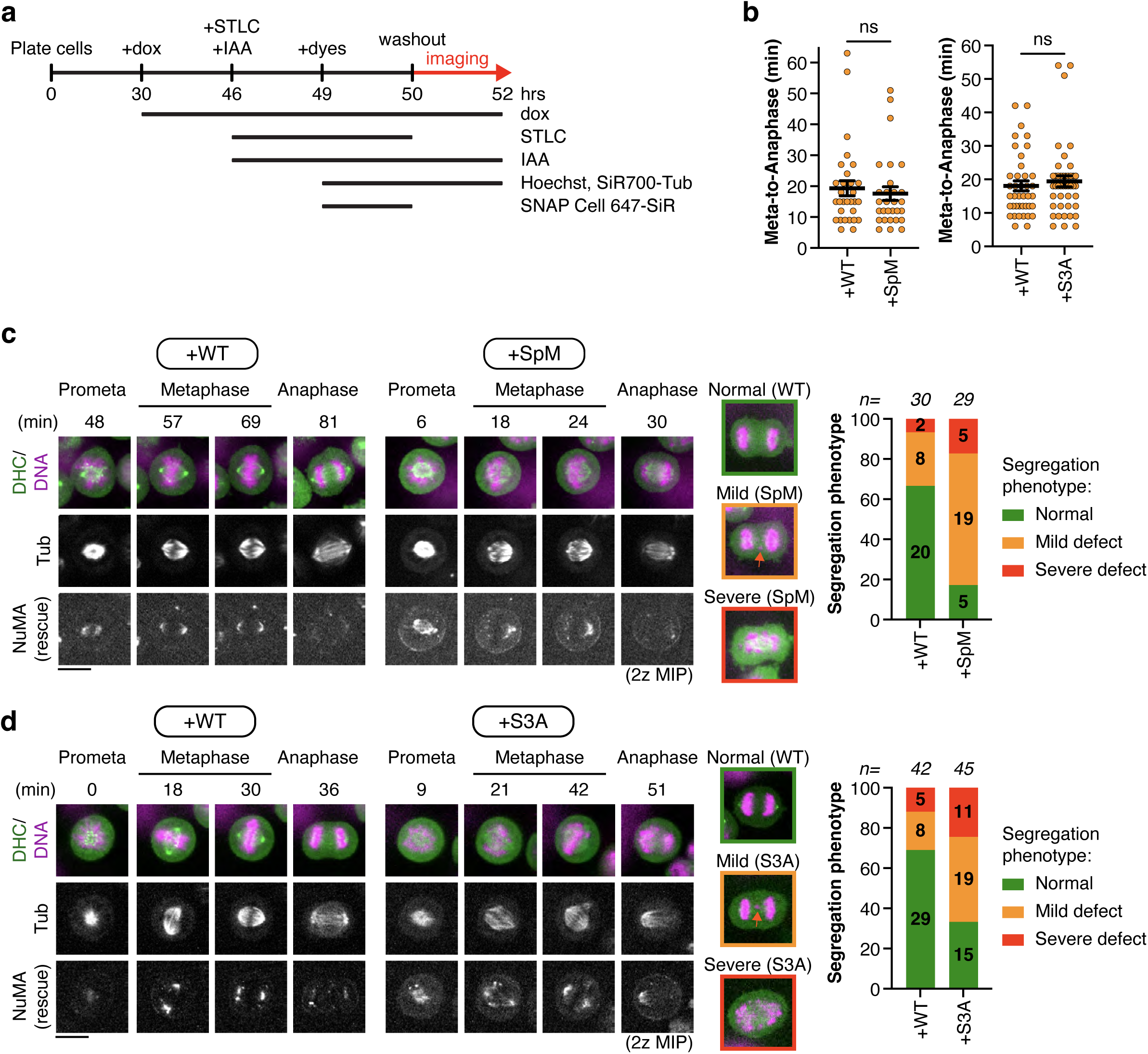
Failure to phosphorylate NuMA’s N-terminus causes chromosome mis-segregation. **a)** Outline of the experimental procedure utilized to express mCherry-NuMA constructs and degrade endogenous NuMA-mACF during prometaphase, followed by STLC-washout and live cell imaging of mitotic progression. **b)** Quantification of time elapsed from spindle bipolarization until anaphase entry after HCT116 cells in which endogenous NuMA-mACF was replaced with mCherry-NuMA WT, SpM or S3A recovered from STLC-washout. n > 29 cells from at least 4 independent experiments; unpaired Welch’s t-tests. **c-d)** Left: Representative live cell fluorescent images showing mitotic progression after STLC-washout in HCT116 cells in which endogenous NuMA-mACF was replaced with either WT, Spindly-like motif mutant (SpM, **c**) or phospho-mutant (S3A, **d**) mCherry-NuMA. Right: Percentages and examples of cells showing either normal, mildly defective (1-2 lagging chromosome or chromosome bridges), or severely defective (multiple errors) chromosome segregation phenotypes. Scale bars represent 10 μm.

## Discussion

Faithful chromosome segregation relies on the concerted action of many spindle-regulating factors, including the microtubule-based dynein adaptor, NuMA^2,3,6^. Despite recent advances in understanding activation and functionality of NuMA-dependent dynein motility^18,19^, cell cycle-regulated assembly and disassembly of the mitotic dynein-dynactin-NuMA (DDN) complex in living cells has remained poorly understood. In this study, we found that CDK1-dependent mitotic phosphorylation of NuMA’s N-terminus regulates its association with dynein/dynactin. This phosphorylation is crucial for dynein function and maintenance of genome stability during mitosis.

The molecular basis of NuMA-dependent dynein activation has long been debated, particularly in light of the multiple conserved dynein-interacting motifs present in its N-terminus and potential autoinhibition by its C-terminus^25,62^. In our cellular system, NuMA-N(1-705) exhibited a robust mitosis-specific recruitment of dynein, indicating that the interaction between NuMA-N and dynein-dynactin is tightly controlled by the cell cycle **(Fig. 1c-f**). Strikingly, expression of membrane-targeted NuMA-N, but not Spindly-N, also consistently impaired chromosome alignment and induced prometaphase arrest **(Fig. 1e, g)**. This phenotype likely results from stronger dynein sequestration to the membrane, which interferes with dynein function at the spindle and kinetochores, since similar phenotypes were observed following DHC depletion^63,64^ or inhibition^13,65^.

Importantly, we reveal that mitotic NuMA phosphorylation at S200 and S203 promotes NuMA binding to dynein **(Fig. 2e, 4a-b, 6c-d, g)**, and that CDK1 is necessary and sufficient for these modifications **(Fig. 3a-b, 4f-g).** At the onset of mitosis, CDK1 directly phosphorylates NuMA S203 near HBS1 **(Fig. 4d, 8a-b)**, which may stabilize NuMA-DHC interaction and/or aid in recruitment of a second dynein motor^31^. We hypothesize that CDK1-Cyclin B1 recruitment to NuMA may involve one of the three conserved Cyclin-binding LxF motifs^66^ at NuMA’s N-terminus **(Fig S7)**. Notably, the *C. elegans* NuMA ortholog, LIN-5, similarly requires CDK1-mediated phosphorylation of its N-terminus to associate with and focus dynein on meiotic spindles^67^. Moreover, CDK1 also phosphorylates T2055 in NuMA’s C-terminal region^36^, which inhibits cortical NuMA accumulation before anaphase^68,69^ and thereby enables the majority of NuMA to interact with spindle microtubules **(Fig. 8a).** In this manner, CDK1 ultimately controls both functionality and localization of DDN through dual phosphorylation of the N- and C-termini of NuMA. In addition to CDK1, PLK1 inhibition also destabilized the DDN complex **(Fig 3a-d, g).** However, considering that the rate of complex disassembly was significantly slower after PLK1 inhibition, we favor a model in which PLK1 is indirectly involved in DDN formation, likely through stimulating CDK1 activity^70,71^. Such a mechanism of CDK1-dependent DDN assembly would be mechanistically distinct from the sequential activation of BICD2, which involves direct phosphorylation by both CDK1 and PLK1^33^. Moreover, while Spindly is also regulated by phosphorylation^30,72^, the mitosis-specific interaction between dynein-dynactin and Spindly-N **(Fig. 1c-f)** was unaffected by short-term CDK1 inhibition **(Fig. S3f-g)**. These findings suggest that while phosphorylation is a common regulatory mechanism for dynein adaptors, the mechanical contribution of phosphorylation and its responsible kinase differs among adaptors. Most dynein adaptors rely on multivalent interactions involving distinct dynein- and dynactin-binding domains^25^. Importantly, dual mutation of both phosphorylation and Spindly-like motifs additively impaired dynein-binding **(Fig. 5a-c**, **Fig 8c).** This is likely explained by the Spindly-like motif (dynactin pointed-end, DCTN5/p25) and HBS1-adjacent phosphorylation sites (DHC) regulating association with distinct parts of dynein-dynactin **(Fig. 8b)**, similar to cryo-EM observations for BICDR1^73^ and consistent with those for NuMA^18^. This multi-site binding model is also supported by findings that NuMA-N (1-705), but neither NuMA-N (1-413) nor NuMA-N (214-705) are sufficient for cortical dynein recruitment in light-inducible targeting assays^13^. Using AID-mediated replacement assays, we confirmed that both CDK1-dependent phosphorylation of NuMA’s N-terminus and the Spindly-like motif are essential for genome stability **(Fig. 6a-b).** Accordingly, replacing NuMA WT with NuMA S3A or NuMA SpM disrupted proper dynein distribution along the spindle, spindle orientation, and chromosome segregation **(Fig. 6c-f an 6c).** Interestingly, however, although both NuMA S3A and NuMA SpM showed reduced dynein binding, micronucleus formation was partially rescued even after mutating both domains **(Fig. 6a-b)**. These results are consistent with recent findings indicating that NuMA’s dynein-independent microtubule crosslinking^74^ activity provides partial spindle stabilization, and underscore NuMA’s multifaceted contributions to mitotic fidelity.

**Figure 8:**
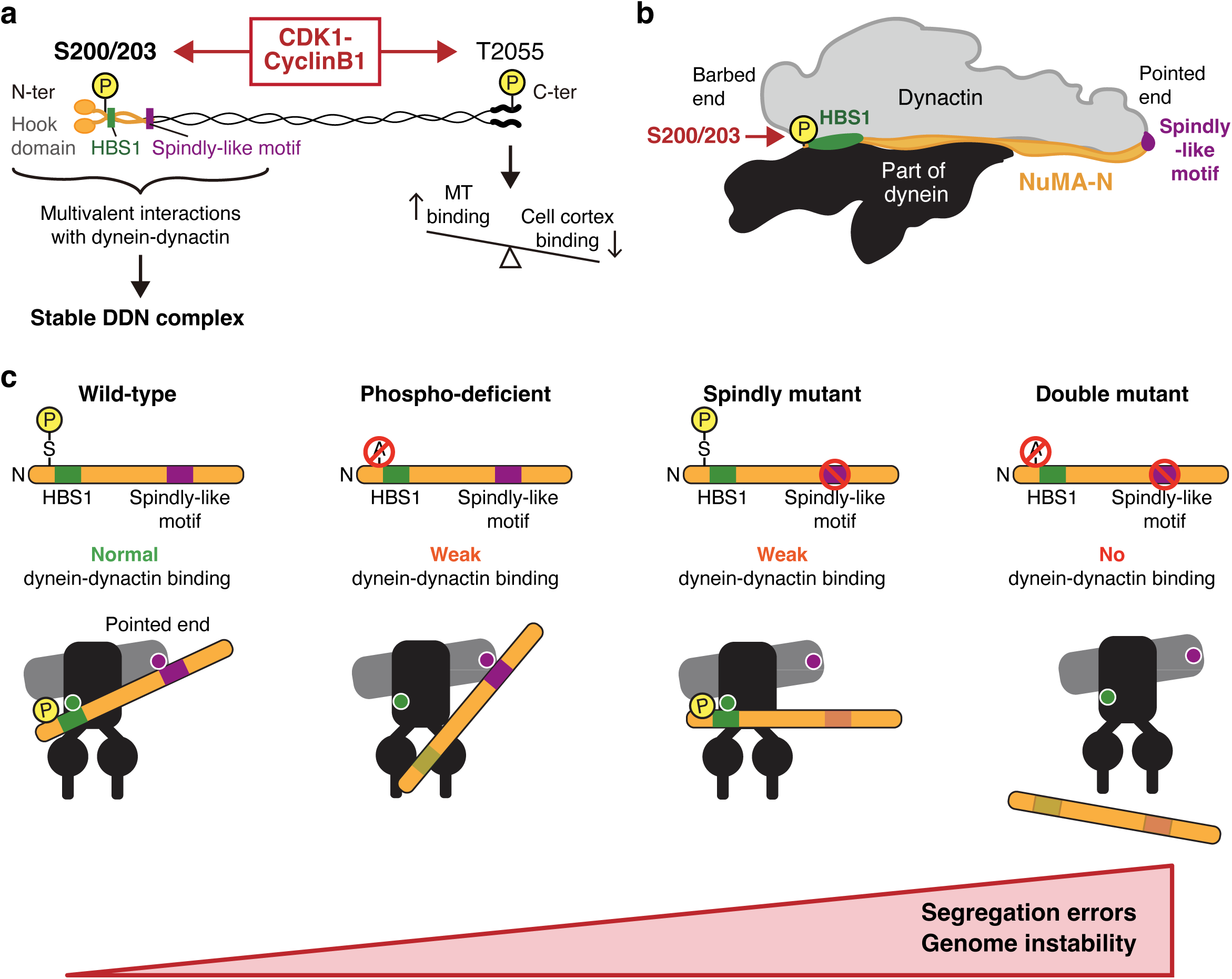
Model of CDK1-dependent NuMA phosphorylation for DDN complex stabilization. **a)** Schematic illustrating CDK1-dependent NuMA phospho-regulation. N-terminal phosphorylation at S200/203 regulates dynein association, whereas C-terminal phosphorylation at T2055 balances spindle and cortical localization. **b)** Model showing multivalent interactions between dynein-dynactin and NuMA. **c)** Diagram depicting how N-terminal NuMA phosphorylation and the Spindly-like motif cooperatively regulate dynein-dynactin binding and preserve genome stability during mitotic cell division.

During anaphase, CDK1 activity decreases, while counteracting PPP2CA phosphatase activity increases sharply^60,75,76^. CDK1 inhibition rapidly dissociates both NuMA-N and full-length NuMA from dynein-dynactin **(Fig. 3f and h),** but considering anaphase’s short duration^77^ (∼10 min), remaining DDN complexes are likely still functional at the anaphase cell cortex. It is tempting to speculate that NuMA’s N-terminal phosphorylation persists for longer than its C-terminal T2055 phosphorylation, because the N-terminal S203 is more sterically hindered from phosphatase access in the DDN complex^18^. Additionally, faster C-terminal NuMA dephosphorylation could promote redistribution of active DDN complexes from the spindle to the anaphase cell cortex **(Fig. 8a)** to pull on astral microtubules^77^. In our system, a phosphomimetic NuMA mutant showed reduced sensitivity to CDK1 inhibition in dynein association assays **(Fig. 4h-i),** but failed to stably recruit dynein during interphase **(Fig. S4f)**, indicating that NuMA phosphorylation alone is insufficient for DDN complex formation in cells. We hypothesize that additional phospho-regulation of dynein and dynactin^78–81^ **(Fig. 2g)** contributes to mitotic DDN assembly and disassembly. Further clarifying the mitotic phospho-regulation of dynein-dynactin-adaptor complexes would provide valuable insights into the relationship between dysregulation of mitotic kinases and cancer^82^.

In conclusion, this study demonstrates a crucial role for N-terminal NuMA phosphorylation and its cooperation with the Spindly-like motif in orchestrating mitotic DDN complex assembly and function in human cells. These results highlight the importance of subtle and multifaceted reversible post-translation modifications in ensuring genome stability during mitosis.

## Acknowledgements

We thank Momoko Nishina, Masako Okumura, Euikyung Yu and Tolkyn Seterkhan for technical assistance and Yoshitoshi Hirao (Instrumental Analysis Section, Core Facilities, Okinawa Institute of Science and Technology Graduate University) for valuable support with LC-MS/MS data acquisition, analyses and advice on sample preparation. GSK461364 was received as a gift from Franz Meitinger (Okinawa Institute of Science and Technology Graduate University). This work was supported by grants from JSPS KAKENHI (21H02481 to TK, 23KF0285 to MvT) and the Okinawa Institute of Science and Technology Graduate University, Japan.

## Author contributions

Conceptualization: TK; Investigation: MvT and TK; Formal analysis: MvT and TK; Resource generation: MvT, KS and TK; Methodology: MvT and TK; Writing: MvT and TK; Supervision: TK; Funding acquisition: TK and MvT

## Methods

### Cell culture, cell line generation and treatments

HCT116 cell lines were cultured in McCoy’s modified 5A medium (Gibco) supplemented with 10% fetal bovine serum (FBS, Gibco) and 1% penicillin-streptomycin (Wako). Flp-In T-REx 293 cells (Invitrogen) were cultured in Dulbecco’s modified Eagle medium (DMEM, Wako), supplemented with 10% FBS and 1% penicillin-streptomycin. All cells were maintained in humidified incubators at 37 °C and 5% CO_2_.

Cell lines were generated by transfection with Effectene (Qiagen), according to the manufacturer’s protocol. 48 h after transfection, cells were selected with 800 μg/mL Neomycin (Roche) or 8 μg/mL Blasticidin S hydrochloride (Funakoshi Biotech) or 200 μg/mL Hygromycin B (Wako) for 12-14 days as described previously^13^. Selected cells were subsequently grouped and validated for expression of integrated constructs in bulk (Flp-In T-REx 293) or isolated as single-cell colonies (HCT116). Single-cell colonies were lysed using DirectPCR lysis solution (Viagen Biotech) according to the manufacturer’s protocol and isolated DNA was PCR-screened for correct insertion of fusion cassettes using Tks Gflex DNA Polymerase (Takara) and primers listed in **Table S6**. All cells were subsequently confirmed mycoplasma-free using a MycoAlert Mycoplasma Detection Kit (Lonza). doxycycline (Milli-Q, Sigma), indole-3-acetic acid (IAA, EtOH, Wako), nocodazole (DMSO, Sigma), BI2536 (DMSO, SelleckChem), MLN8237 (DMSO, SelleckChem), RO3306 (DMSO, Sigma), MG132 (DMSO, Sigma), GSK461364 (DMSO, SelleckChem), D4476 (DMSO, Funakoshi), proTAME (DMSO, R&D Systems), apcin (DMSO, Cayman Chemicals) and STLC (DMSO, Sigma) stocks were prepared in their appropriate solvents and added to culture media at indicated times. Established human tissue culture cell lines, as well as PCR primers and guideRNAs used to generate and confirm genotypes of cell lines in this study are summarized in **Tables S4, S6 and S7**, respectively.

### Plasmid construction

Plasmids for CRISPR/Cas9-mediated genome editing and auxin-inducible degron methodology were constructed as described previously^13,16^. To make plasmid constructs for the membrane-tethering assay, MluI site of pcDNA5FRT/TO plasmid was mutated, and then a NheI site was introduced between HindIII and KpnI sites by PCR-based site-directed mutagenesis (pTK764). Subsequently, an mCherry-3xFlag-CAAX fragment amplified by PCR was inserted between the NheI and XhoI sites (pTK887). To create Adaptor-mCherry-3xFlag-CAAX plasmids, adaptor fragments were inserted in the MluI site of pTK887 for NuMA-N (pTK888), NuMA-N SpM (pTK889), Hook3-N(pTK951), Hook3-N I152A (pTK952), BICD2-N (pTK953) and Spindly-N (pMT20).

PCR-based site-directed mutagenesis to generate phosphomutants was carried out using PrimeSTAR Max DNA Polymerase (Takara), according to the manufacturer’s protocol. PCR primers used for site-directed mutagenesis are included in **Table S6.** Mutations were confirmed by Sanger sequencing.

### Hypotonic immunoprecipitation

Cells from one to three near-confluent 15-cm plates and collected by trypsinization, washed twice with ice-cold PBS (Shimadzu Diagnostics) and lysed with freshly prepared hypotonic lysis buffer (50 mM Tris-HCl pH7.5, 10 mM NaCl, 2.5 mM MgCl_2_, 1 mM DTT and 0.01% digitonin (TCI) in Milli-Q water with EDTA-free protease and phosphatase inhibitors and 50 U/mL Benzonase® nuclease (Merck) added freshly). Cell lysates were incubated for 1 hr with constant rotation at 4 °C before being cleared by centrifugation (13200 rpm for 15 min at 4 °C). Cleared lysates were incubated with 30-40 μL pre-equilibrated Flag M2 agarose (Sigma), GFP-Trap agarose (ChromoTek) or RFP-Trap agarose (ChromoTek) beads for 2 h with constant rotation. Beads were pelleted by centrifugation (700 rcf for 2 min at 4 °C), washed 5 times with hypotonic lysis buffer and proteins eluted with 2× sample buffer (100 mM Tris-HCl pH6.8, 4% SDS and 10% glycerol) for 5 min at 98 °C and processed for either immunoblotting or mass spectrometry. 0.004% Bromophenol blue and 2.5% β-mercaptoethanol were added to sample buffer for samples destined for immunoblotting.

### Crosslinked immunoprecipitation

Crosslinked immunoprecipitation was carried out as described previously^83^, with minor adjustments. Cells from two to four near-confluent 15-cm plates were treated as described and collected by trypsinization. Cells were washed twice with serum-free DMEM and crosslinked with 1 mM dithiobis(succinimidyl propionate) in serum-free DMEM for 30 min with constant movement, followed by quenching with a final concentration of 25 mM Tris-HCl pH7.5 for 10 min. Cells were pelleted by centrifugation (1500 rpm for 3 min at 4 °C), washed twice with PBS and lysed in RIPA buffer (50 mM Tris-HCl pH7.5, 150 mM NaCl, 1% NP-40 substitute, 0.5% sodium deoxycholate and 0.1% SDS in Milli-Q). 50 U/mL Benzonase, EDTA-free protease and phosphatase inhibitors were also added freshly. After 30 min, lysates were cleared by centrifugation (13200 rpm for 15 min at 4 °C) and supernatants were incubated with 30-40 μL pre-equilibrated Flag M2 agarose or GFP-Trap agarose beads for 2 h with constant rotation. Beads were spun down (700 rcf for 2 min at 4 °C) and washed with RIPA buffer 5 times before proteins were eluted with 2× sample buffer (100 mM Tris-HCl pH6.8, 4% SDS and 10% glycerol) for 5 min at 98 °C before being processed for immunoblotting or mass spectrometry.

### Biotin-based proximity labeling

NuMA-mClover-miniTurbo knock-in cells were grown in DMEM supplemented with 10% FBS and 1% penicillin-streptomycin to reduce background biotinylation. Two to three 15-cm plates were prepared as described and 50 μM biotin (TCI) was added to the cells, followed by 30-min incubation in the incubator. Afterward, cells were collected by trypsinization and washed 6 times with ice-cold PBS to remove excess biotin. Cells were subsequently lysed with RIPA buffer for 30 min and cleared by centrifugation (13200 rpm for 15 min at 4 °C). Supernatants were collected and incubated with pre-equilibrated Pierce streptavidin-agarose resin (ThermoFisher) for 2 h with constant rotation. Beads were collected by centrifugation, washed 5 times with RIPA buffer and eluted with 2× sample buffer for 5 min at 98 °C. The experiment in **Fig. S2f** was performed using a 6-well plate to prepare the lysates and without performing immunoprecipitation.

### Immunoblotting

Cell lysates or immunoprecipitates were resolved by SDS-PAGE using 7.5% or 10% TGX FastCast acrylamide gels (Bio-Rad), cast according to the manufacturer’s guidelines. Proteins were separated using running buffer (25 mM TRIS, 190 mM Glycine and 0.1% SDS) and transferred to methanol-activated Immobilon PVDF membranes (0.45 μm, Merck) overnight at 30 mA in transfer buffer (0.025 M Tris, 0.19 M Glycine and 10% ethanol). Membranes were blocked for 1 h with 5% skim milk powder (Wako) in PBS and subsequently incubated with the indicated primary antibodies **(Table S5)**, diluted in PBS for 2 h with constant shaking. Thereafter, membranes were washed 5 times with 0.05% Tween-20 in PBS and incubated with the appropriate HRP-conjugated secondary antibodies (Cytiva, **Table S5**) for 1 h with constant shaking. Membranes were subsequently washed an additional 5 times with 0.05% Tween-20 in PBS and developed with either ECL Western Blotting Detection Reagent (Cytiva) or ECL Prime Western Blotting Detection Reagent (Cytiva), according to the manufacturer’s protocol. Membranes were imaged using a Fusion Solo S imaging system (Vilber).

### Mass spectrometry and data analysis

Mass spectrometry samples were prepared with S-Trap spin columns (Protifi), according to the manufacturer’s protocol. Briefly, eluted proteins were reduced with 5 mM TCEP (15 min at 37 °C, TCI), thiol-blocked with 20 mM MMTS (15 min at 37 °C, TCI) and denatured with a final concentration of ∼4.5% phosphoric acid (Wako) before being bound with S-Trap binding buffer (100 mM TEAB pH7.55 in 90% methanol). Samples were loaded onto S-Trap columns, spun down for 2 min at 4000 rpm, washed 5 times with S-Trap binding buffer and subsequently incubated with 2 μg Trypsin Platinum (Promega, diluted in 50 mM TEAB) for 2h at 47 °C using a water bath. Digested peptides were eluted with elution buffers 1 (50 mM TEAB), 2 (0.2% formic acid) and 2(50% acetonitrile) and lyophilized using a Scientific EZ-2 Elite Evaporator (SP Genevac) for 2 h at 42 °C using the Aqueous setting. Peptides were then dissolved in 0.1% formic acid and concentrations were determined using a NanoDrop One^C^ (ThermoFisher).

Peptides were diluted to 100 ng/μL, and 2.5 μL duplicates or triplicates of each sample were analyzed by LC-MS using an Orbitrap Fusion Lumos (ThermoFisher) in data-dependent acquisition mode coupled to an ACQUITY UPLC M-Class nanoflow liquid chromatography system (Waters). Peptides were separated on trap (2 cm x 180 μm nanoEase M/Z Symmetry C18 Trap Column) and separation (15 cm x 75 μm nanoEase M/Z HSS T3 Column) columns at a flow rate of 500 nL/min with a multistep gradient. Mobile phases used to elute injected peptides from the analytical column were 0.1% formic acid in water (A) and 0.1% formic acid in acetonitrile (B), with a linear gradient from 5% B to 30% B in 60 min. The mass spectrometer was set to perform data acquisition in the positive ion mode with a scan range of 380-1,500 m/z for MS and auto for MS/MS. The cycle time was set to 3 sec and charge states 2-7 were selected from each survey scan and subjected to HCD fragmentation. MS and MS/MS scan parameter settings were as follows: collision energy, 30%; electrospray voltage, 2.0 kV; capillary temperature, 275℃.

MS and MS/MS data were analyzed using Proteome Discoverer v3.2 (ThermoFisher) with either the CHIMERYS (for label-free quantification) or SequestHT-based (for additional modification discovery) algorithms. Data were queried against a UniProtKB/SWISS-Prot database (v2024-03; *Homo sapiens*; 20360 sequences). All database searches were performed using a precursor mass tolerance of ± 10 ppm and fragment ion mass tolerance of ± 0.6 Da (SequestHT) or fragment mass tolerance of ± 0.3 Da (CHIMERYS) Enzyme specificity was set to trypsin and maximum missed cleavages was set to 2. Cysteine carbamidomethylation was set as a fixed modification, while phosphorylation (S/T/Y) was set as a dynamic modification. Unique and razor peptides were quantified based on intensity and normalized to total peptide amount. Common and cell culture-specific contaminants were discarded^84^. Percolator was used to adjust the false discovery rate (FDR) to 1% at the peptide level.

### Live cell imaging

Live cell imaging was performed using an ECLIPSE Ti-E inverted spinning disk confocal microscope with perfect focus system (Nikon) that was connected to a CSU-W1 confocal scanner unit (Yokogawa Electric Corporation) and an ORCA-Flash 4.0 digital CMOS camera (Hamamatsu Photonics). The microscope was equipped with a 60× 1.40 numerical aperture objective lens (Plan Apo λ, Nikon), 488-, 561-, 640- and 685-nm lasers (Coherent) for a SOLA LED light engine (Lumencor) to visualize DNA from cells stained with Hoechst® 33342 (Sigma).

Live cell confocal imaging was basically carried out as described before^11,13,16^, with minor adjustments. All live cell imaging experiments were carried out in a stage-top incubator (Tokai Hit) maintained at 37°C and 5% CO_2_. Cells were seeded on glass-bottom dishes (CELLview™, #627860 or #627870, Greiner Bio-One) and DNA, SNAP-tagged proteins and microtubules were pre-stained with 50 ng/mL Hoechst® 33342 (Sigma), 120 nM SNAP Cell 647-SiR and/or 50 nM SiR700-tubulin (Spirochrome), respectively, for >30 min before observation where indicated.

For snap shots, five or more z-section images with 1-μm spacing were acquired. Maximum intensity projections of select slices or single z-section images are shown as indicated in each figure. Signals were linearly adjusted using Fiji and Photoshop to optimize image clarity. To obtain the orthogonal (x-z) images shown in **Fig. 5c**, at least 31 z-section images with 0.2 μm spacing were obtained for each cell.

Time-lapse imaging with NuMA-N(1-705)-mCherry-3xFlag-CAAX-expressing Flp-In T-REx 293 DHC-mClover KI cells was carried out by addition of 2 μg/mL Dox overnight and 50 ng/mL Hoechst® 33342 1 h before imaging. Image acquisition was initiated directly after addition of the indicated drugs, three z-section images with 1-μm spacing were acquired every 3 min for 2 h.

For STLC washout experiments, 2 μg/mL Dox was added to asynchronously growing cells overnight. The next day, 10 μM STLC and 500 μM IAA were added for 4 h to accumulate prometaphase cells, with DNA, SNAP, and tubulin dyes additionally added during the last hour. Cells were released from prometaphase arrest by washing cells in pre-warmed culture medium for 3-5 times, after which they were incubated with medium containing 2 μg/mL Dox, IAA and DNA and tubulin dyes and moved to the microscope stage. Two z-section images with 1-μm spacing were then acquired every 3 min for 2 h. STLC washout data are shown as 2z maximum intensity projections.

### Immunoprecipitation of CAAX-conjugated constructs and *in vitro* phosphorylation

Two near-confluent 15-cm plates were treated with 2 μg/mL Dox overnight (with 9 μM RO3306 or 10 μM STLC additionally added where indicated) and collected by trypsinization, followed by washing twice with ice-cold PBS. Cells were lysed in IP buffer (50 mM Tris-HCl pH7.5, 100 mM KCl, 1 mM MgCl_2_, 0.5% NP-40 substitute, 0.1% sodium deoxycholate and 0.01% digitonin with 50 U/mL Benzonase, EDTA-free protease and phosphatase inhibitors freshly added) for 30 min with constant rotation and centrifuged (13200 rpm for 15 min at 4 °C). Cleared supernatants were incubated with 30 μL of pre-equilibrated Flag M2 agarose beads for 2 h with constant rotation, before beads were collected by centrifugation and washed twice with IP buffer (for subsequent *in vitro* phosphorylation) or 5 times (for other experiments).

For *in vitro* phosphorylation experiments, beads were subsequently incubated with 50 μL kinase buffer (20 mM HEPES-KOH pH7.7, 80 mM KCl, 5 mM MgCl_2_ and 2 mM ATP) containing 20 nM modified *H. sapiens* CDK1-Cyclin B1, 1 μM *S. pombe* Suc1 (both described in Shintomi *et al.*^59^) and 9 μM RO3306 where indicated for 15 min at room temperature. Beads were then washed with IP buffer three more times. Bound proteins were eluted using 2× sample buffer for 5 min at 98 °C and processed for mass spectrometry or immunoblotting.

### Signal quantification

Signals were quantified from 16-bit images using ImageJ. Cortical dynein levels in Flp-In T-REx 293 DHC-mClover cells were quantified by dividing the maximum signal along a straight line (line width 3 or 5) spanning the cells by the average cytoplasmic background in a single z-section, after subtraction of the non-cellular background from both signals. Dynein distribution **(Fig. 6d)** was determined by dividing the maximum pole accumulation (determined using a 5×5 pixel box along the spindle pole) by the average spindle signal (determined using a 20×10 pixel using 3z MIPs. Both signals were corrected for non-cellular background signals. Dynein accumulation at the spindle poles **(Fig. 5e)** was determined by subtracting cytoplasmic values from maximum pole accumulation values (5×5 pixel box along the spindle pole).

### Optogenetic targeting

NuMA-N(1-705)-RFP-Nano was conditionally expressed by doxycycline treatment and targeted to membrane in interphase HCT116 cells by local light illumination using a Mosaic 3 device, as described in metaphase cells^13^. Hook3-N(1-552)-RFP-Nano-expressing cells were generated using the same strategy as NuMA-N(1-705)-RFP-Nano^13^.

### Statistics

No statistical methods were used to pre-determine sample sizes. Sample sizes were chosen based on previously published studies and pilot experiments. Significance was calculated using GraphPad Prism 10.4.1 (Dotmatics). Generally, Brown-Forsythe and Welch ANOVA with Dunnett’s multiple comparisons or unpaired Welch’s t-tests were used to calculate significance. p<0.05 was considered statistically significant and p-values are reported as *p<0.05, **p<0.01 and ***p<0.001.

## Supplemental tables

**Table S1:** Mass spectrometry data of crosslinked NuMA immunoprecipitation experiments from asynchronized (Control) or mitotically-enriched (Sample) NuMA-mACF KI HCT116 cells.

**Table S2:** Mass spectrometry data of biotin-based proximity labeling followed by immunoprecipitation of biotinylated proteins from asynchronized (Control) or mitotically-enriched (Sample) NuMA-mClover-miniTurbo KI HCT116 cells.

**Table S3:** Mass spectrometry data of crosslinked DCTN2 immunoprecipitation experiments from asynchronized (Control) or mitotically-enriched (Sample) DCTN2-mACF KI HCT116 cells.

**Table S4:** Cell lines used in this study.

| No | Name | Description | Clone | Plasmids | Parent | Reference |
| --- | --- | --- | --- | --- | --- | --- |
| 1 | Flp-In T-REx 293 + DHC-mClover | *DYNC1H1*:: DHC-mClover (Neo) | No. 12 | pTK308 + pTK345 | Flp-In T-Rex 293 (Invitrogen)<sup>45</sup> | This study |
| 2 | Flp-In T-REx 293 + DHC-mClover + BICD2-N(1-400)-mCh-CAAX | *DYNC1H1*:: DHC-mClover (Neo) FRT site:: BICD2-N(1-400)-mCherry-3xFlag-CAAX (Hyg) | N/A | pOG44 + pTK953 | 1 | This study |
| 3 | Flp-In T-REx 293 + DHC-mClover + Hook3-N(1-552)-mCh-CAAX | *DYNC1H1*:: DHC-mClover (Neo) FRT site:: Hook3-N(1-552)-mCherry-3xFlag-CAAX (Hyg) | N/A | pOG44 + pTK951 | 1 | This study |
| 4 | Flp-In T-REx 293 + DHC-mClover + Hook3-N(1-552, I154A)-mCh-CAAX | *DYNC1H1*:: DHC-mClover (Neo) FRT site:: Hook3-N(1-552, I154A)-mCherry-3xFlag-CAAX (Hyg) | N/A | pOG44 + pTK952 | 1 | This study |
| 5 | Flp-In T-REx 293 + DHC-mClover + NuMA-N(1-705)-mCh-CAAX | *DYNC1H1*:: DHC-mClover (Neo) FRT site:: NuMA-N(1-705)-mCherry-3xFlag-CAAX (Hyg) | N/A | pOG44 + pTK888 | 1 | This study |
| 6 | Flp-In T-REx 293 + DHC-mClover + Spindly-N(1-359)-mCh-CAAX | *DYNC1H1*:: DHC-mClover (Neo) FRT site:: Spindly-N(1-359)-mCherry-3xFlag-CAAX (Hyg) | N/A | pOG44 + pMT20 | 1 | This study |
| 7 | HCT116 *tet*-OsTIR1 DCTN2-mACF | *AAVS1*:: PTRE3G OsTIR1 (Puro), *DCTN2*:: DCTN2-mAID-mClover-3xFlag (Neo) | No. 7 | pTK950 + pTK493 | HCT116 *tet*-OsTIR1<sup>63</sup> | This study |
| 8 | HCT116 *tet*-OsTIR1 DCTN2-mACF + DCTN1-SNAP | *AAVS1*:: PTRE3G OsTIR1 (Puro), *DCTN2*:: DCTN2-mAID-mClover-3xFlag (Neo), *DCTN1*:: DCTN1-SNAP (Blast) | No.17 | pTK525 + pTK672 | 7 | This study |
| 9 | HCT116 *tet*-OsTIR1 NuMA-mACF + SNAP-LGN | *AAVS1*:: PTRE3G OsTIR1 (Puro), *NuMA1*:: NuMA-mAID-mClover-3xFlag (Neo), *GPSM2*:: SNAP-LGN (BSD) | No. 3 | pTK473 + pTK586 | HCT116 *tet*-OsTIR1 NuMA-mACF #1 | Okumura *et al.*<sup>13</sup> |
| 10 | Flp-In T-REx 293 + DHC-mClover + NuMA-N(1-705, T107A/S109A)-mCh-CAAX | *DYNC1H1*:: DHC-mClover (Neo) FRT site:: NuMA-N(1- | N/A | pOG44 + pMT34 | 1 | This study |
|  |  | 705, T107A/S109A)-mCherry-3xFlag-CAAX (Hyg) |  |  |  |  |
| 11 | Flp-In T-REx 293 + DHC-mClover + NuMA-N(1-705, S189A/S190A/S191A/S192A)-mCh-CAAX | *DYNC1H1*:: DHC-mClover (Neo) FRT site:: NuMA-N(1-705, S189A/S190A/S191A/S192A)-mCherry-3xFlag-CAAX (Hyg) | N/A | pOG44 + pMT30 | 1 | This study |
| 12 | Flp-In T-REx 293 + DHC-mClover + NuMA-N(1-705, S198A/S200A/S203A, S3A)-mCh-CAAX | *DYNC1H1*:: DHC-mClover (Neo) FRT site:: NuMA-N(1-705, S198A/S200A/S203A, S3A)-mCherry-3xFlag-CAAX (Hyg) | N/A | pOG44 + pMT31 | 1 | This study |
| 13 | Flp-In T-REx 293 + DHC-mClover + NuMA-N(1-705, S271A)-mCh-CAAX | *DYNC1H1*:: DHC-mClover (Neo) FRT site:: NuMA-N(1-705, S271A)-mCherry-3xFlag-CAAX (Hyg) | N/A | pOG44 + pMT49 | 1 | This study |
| 14 | Flp-In T-REx 293 + DHC-mClover + NuMA-N(1-705, A368V/A369V, CC1-likeM)-mCh-CAAX | *DYNC1H1*:: DHC-mClover (Neo) FRT site:: NuMA-N(1-705, A368V/A369V, CC1-likeM)-mCherry-3xFlag-CAAX (Hyg) | N/A | pOG44 + pMT33 | 1 | This study |
| 15 | Flp-In T-REx 293 + DHC-mClover + NuMA-N(1-705, T211A)-mCh-CAAX | *DYNC1H1*:: DHC-mClover (Neo) FRT site:: NuMA-N(1-705, T211A)-mCherry-3xFlag-CAAX (Hyg) | N/A | pOG44 + pMT47 | 1 | This study |
| 16 | Flp-In T-REx 293 + DHC-mClover + NuMA-N(1-705, S3A+T211A)-mCh-CAAX | *DYNC1H1*:: DHC-mClover (Neo) FRT site:: NuMA-N(1-705, S3A+T211A)-mCherry-3xFlag-CAAX (Hyg) | N/A | pOG44 + pMT48 | 1 | This study |
| 17 | Flp-In T-REx 293 + DHC-mClover + mCh-CAAX | *DYNC1H1*:: DHC-mClover (Neo) FRT site:: mCherry-3xFlag-CAAX (Hyg) | N/A | pOG44 + pTK887 | 1 | This study |
| 18 | Flp-In T-REx 293 + DHC-mClover + NuMA-N(1-705, S198D/S200D/S203D, S3D)-mCh-CAAX | *DYNC1H1*:: DHC-mClover (Neo) FRT site:: NuMA-N(1-705, S198D/S200D/S203D, S3D)-mCherry-3xFlag-CAAX (Hyg) | N/A | pOG44 + pMT52 | 1 | This study |
| 19 | Flp-In T-REx 293 + DHC-mClover + NuMA-N(1-705, L413A/E416A/L418A/E421A, SpM)-mCh-CAAX | *DYNC1H1*:: DHC-mClover (Neo) FRT site:: NuMA-N(1-705, L413A/E416A/L418A/E421A, SpM)-mCherry-3xFlag-CAAX (Hyg) | N/A | pOG44 + pTK889 | 1 | This study |
| 20 | Flp-In T-REx 293 + DHC-mClover + NuMA-N(1-705, Q124A/L131A, HookM)-mCh-CAAX | *DYNC1H1*:: DHC-mClover (Neo) FRT site:: NuMA-N(1-705, Q124A/L131A, HookM)-mCherry-3xFlag-CAAX (Hyg) | N/A | pOG44 + pEK1 | 1 | This study |
| 21 | Flp-In T-REx 293 + DHC-mClover + NuMA-N(HookM/SpM)-mCh-CAAX | *DYNC1H1*:: DHC-mClover (Neo) FRT site:: NuMA-N(1-705, HookM/SpM)-mCherry-3xFlag-CAAX (Hyg) | N/A | pOG44 + pEK2 | 1 | This study |
| 22 | Flp-In T-REx 293 + DHC-mClover + NuMA-N(1-705, S3A/SpM)-mCh-CAAX | *DYNC1H1*:: DHC-mClover (Neo) FRT site:: NuMA-N(1- | N/A | pOG44 + pMT53 | 1 | This study |
|  |  | 705, S3A/SpM)-mCherry-3xFlag-CAAX (Hyg) |  |  |  |  |
| 23 | HCT116 *tet*-OsTIR1 NuMA-mACF + DHC-SNAP | AAVS1:: PTRE3G OsTIR1 (Puro), *NuMA1*:: NuMA-mAID-mClover-3xFlag (Neo), *DYNC1H1*:: DHC-SNAP (BSD) | No. 18 | pTK308 + pTK600 | HCT116 *tet*-OsTIR1 NuMA-mACF #1 | Okumura *et al.*<sup>13</sup> |
| 24 | HCT116 *tet*-OsTIR1 NuMA-mACF + DHC-SNAP + mCherry-NuMA (WT) | AAVS1:: PTRE3G OsTIR1 (Puro), *NuMA1*:: NuMA-mAID-mClover-3xFlag (Neo), *DYNC1H1*:: DHC-SNAP (BSD), *Rosa26*:: PTRE3G mCherry-NuMA(WT) (Hygro) | No. 7 | hROSA26 CRISPR-pX330 + pTK503 | 16 | Okumura *et al.*<sup>13</sup> |
| 25 | HCT116 *tet*-OsTIR1 NuMA-mACF + DHC-SNAP + mCherry-NuMA (S2A) | AAVS1:: PTRE3G OsTIR1 (Puro), *NuMA1*:: NuMA-mAID-mClover-3xFlag (Neo), *DYNC1H1*:: DHC-SNAP (BSD), *Rosa26*:: PTRE3G mCherry-NuMA(S2A) (Hygro) | No. 12 | hROSA26 CRISPR-pX330 + pMT54 (#B5) | 16 | This study |
| 26 | HCT116 *tet*-OsTIR1 NuMA-mACF + DHC-SNAP + mCherry-NuMA (S3A) | AAVS1:: PTRE3G OsTIR1 (Puro), *NuMA1*:: NuMA-mAID-mClover-3xFlag (Neo), *DYNC1H1*:: DHC-SNAP (S3A), *Rosa26*:: PTRE3G mCherry-NuMA(WT) (Hygro) | No. 10 | hROSA26 CRISPR-pX330 + pMT54 (#1L2) | 16 | This study |
| 27 | HCT116 *tet*-OsTIR1 NuMA-mACF + DHC-SNAP + mCherry-NuMA (S3D) | AAVS1:: PTRE3G OsTIR1 (Puro), *NuMA1*:: NuMA-mAID-mClover-3xFlag (Neo), *DYNC1H1*:: DHC-SNAP (BSD), *Rosa26*:: PTRE3G mCherry-NuMA(S3D) (Hygro) | No. 5 | hROSA26 CRISPR-pX330 + pMT59 | 16 | This study |
| 28 | HCT116 *tet*-OsTIR1 NuMA-mACF + DHC-SNAP + mCherry-NuMA (SpM) | AAVS1:: PTRE3G OsTIR1 (Puro), *NuMA1*:: NuMA-mAID-mClover-3xFlag (Neo), *DYNC1H1*:: DHC-SNAP (BSD), *Rosa26*:: PTRE3G mCherry-NuMA(SpM) (Hygro) | No. 11 | hROSA26 CRISPR-pX330 + pTK694 | 16 | This study |
| 29 | HCT116 *tet*-OsTIR1 NuMA-mACF + DHC-SNAP + mCherry-NuMA (S3A/SpM) | AAVS1:: PTRE3G OsTIR1 (Puro), *NuMA1*:: NuMA-mAID-mClover-3xFlag (Neo), *DYNC1H1*:: DHC-SNAP (BSD), *Rosa26*:: PTRE3G mCherry-NuMA(S3A/SpM) (Hygro) | No. 11 | hROSA26 CRISPR-pX330 + pMT55 | 16 | This study |
| 30 | DHC-SNAP+NuMA(1-705)-RFP-Nano | AAVS1::PCMV Mem-BFP-iLID (Puro), *DYNC1H1*::DHC-SNAP (BSD), *Rosa26*:: PTRE3G NuMA(1-705)-RFP-Nano (Hygro) | No.5 | hROSA26 CRISPR-pX330 +pTK631 | Mem-BFP-iLID+DHC-SNAP #11 | Okumura *et al.*<sup>13</sup> |
| 31 | DHC-SNAP+Hook3(1-552)-RFP-Nano | AAVS1::PCMV Mem-BFP-iLID (Puro), *DYNC1H1*::DHC-SNAP (BSD), *Rosa26*:: PTRE3G Hook3(1-552)-RFP-Nano (Hygro) | No. 11 | hROSA26 CRISPR-pX330 +pTK633 | Mem-BFP-iLID+DHC-SNAP #11 | This study |

**Table S5:** Antibodies list and working dilutions.

| Antibody | Origin | Source | Dilution |
| --- | --- | --- | --- |
| Anti-NuMA | Rabbit | Abcam, ab36999 | 1/2000 |
| Anti-DCTN1/p150 | Mouse | BD Biosciences, 610473 | 1/2000 |
| Anti-DHC | Rabbit | Santa Cruz, sc-9115 | 1/500 |
| Anti-Flag (M2) | Mouse | Sigma, F3165 | 1/2000 |
| Anti-Hook3 | Rabbit | Proteintech, 15457-1-AP | 1/1000 |
| Anti-LGN | Rabbit | Bethyl, A303-032A | 1/1000 |
| Anti-Rabbit, HRP-linked | Donkey | Cytiva, NA934 | 1/10000 |
| Anti-Mouse, HRP-linked | Sheep | Cytiva, NA931 | 1/10000 |

**Table S6:**
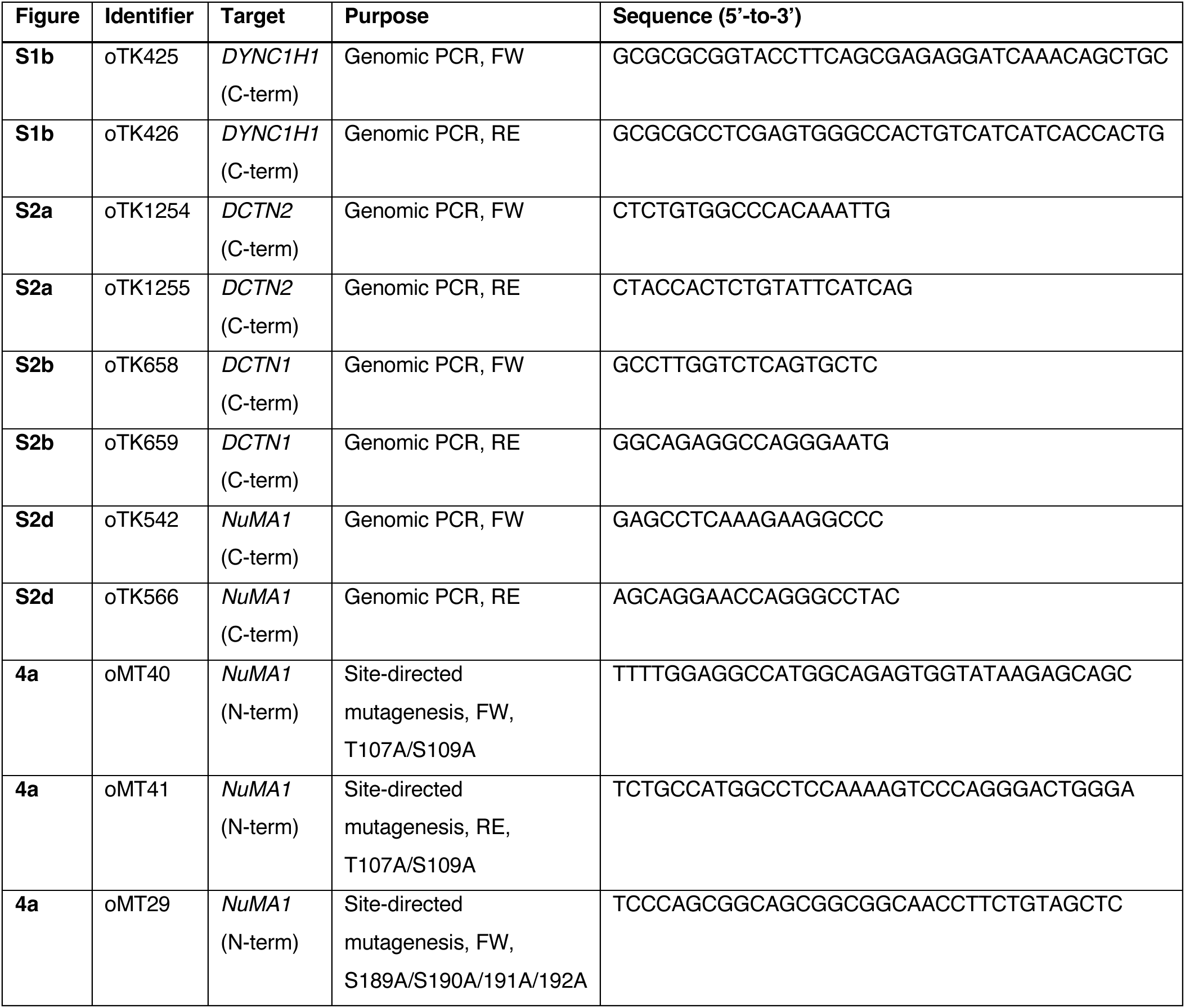
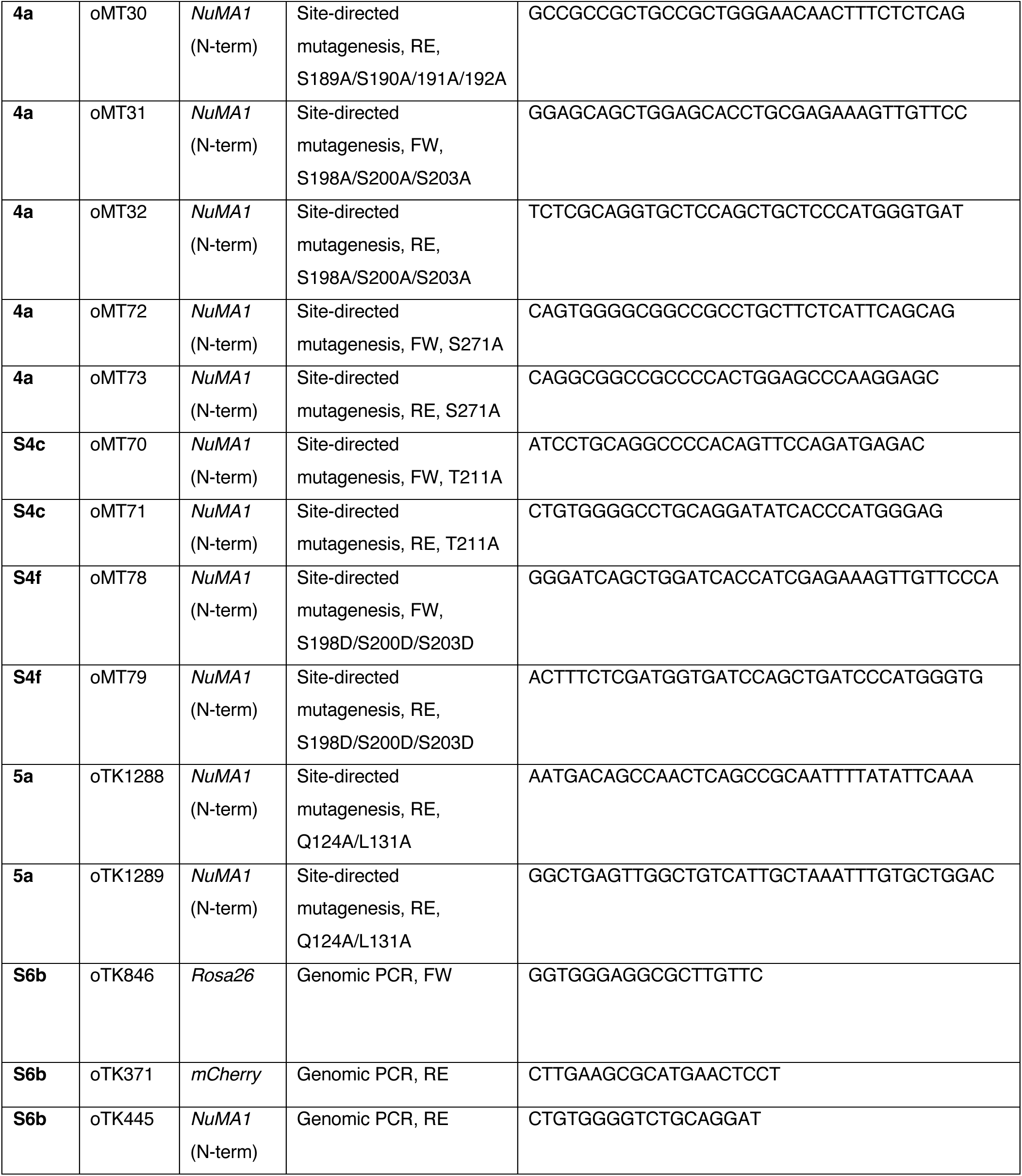
Oligonucleotides used in this study.

**Table S7:**
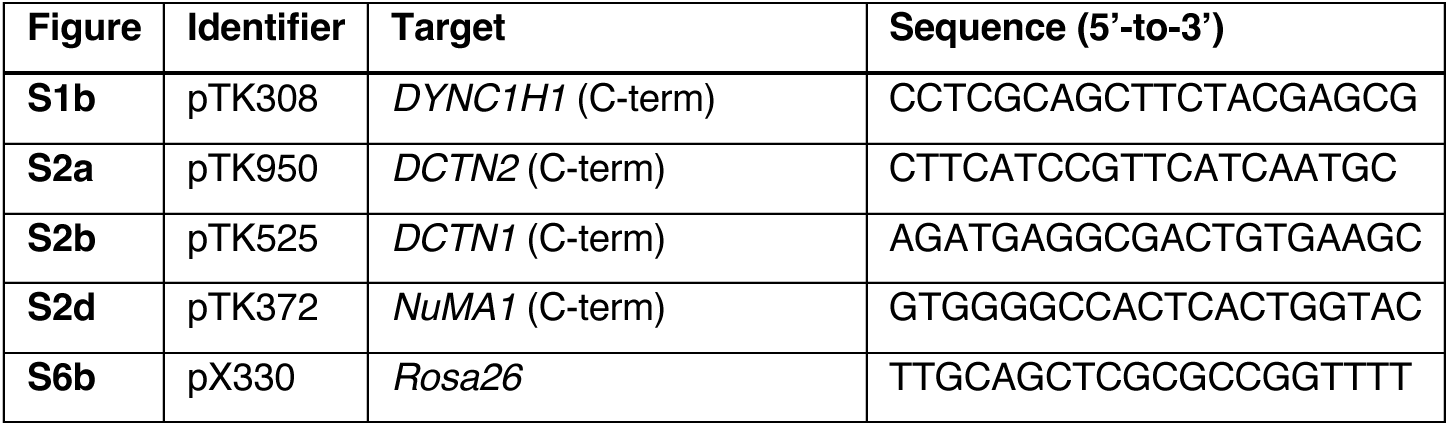
sgRNAs used in this study.

## Supplementary figure legends

**Figure S1:**
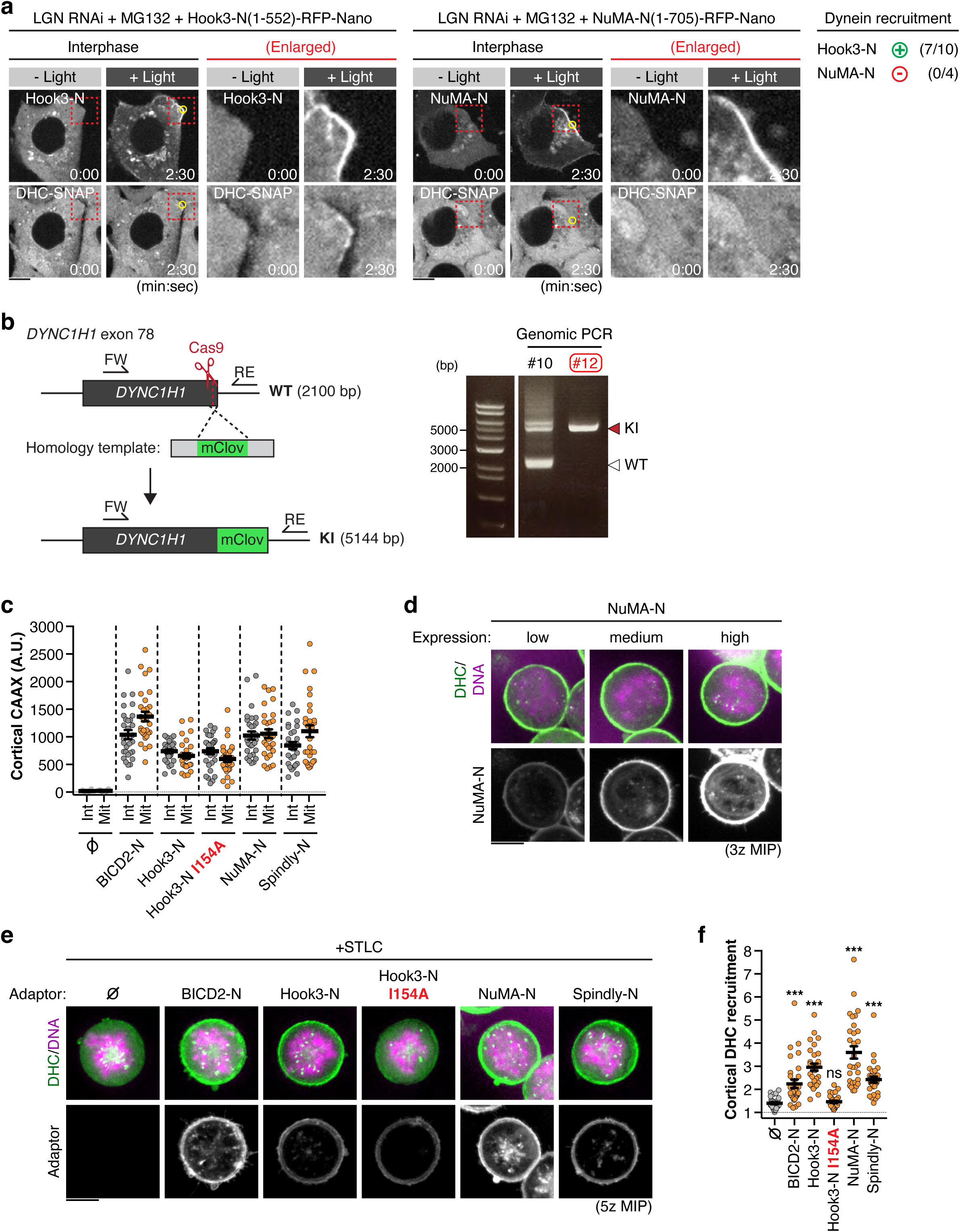
Membrane tethering of dynein adaptor fragments. **a)** Live fluorescent images showing DHC-SNAP recruitment to the cell membrane in interphase HCT116 cells expressing Mem-BFP-iLID and either Hook3-N(1-552)-RFP-Nano or NuMA-N(1-705)-RFP-Nano. Cells were LGN-depleted and MG132-treated before imaging to compare DHC-SNAP recruitment between interphase and metaphase cells. Yellow circles indicate the light-illuminated region. **b)** Left: Schematic representation of the generation of DHC-mClover Flp-In T-REx 293 KI cells. The mClover cassette (including a Neomycin selection marker depicted as mClover for simplicity), was inserted at the C-terminus of the *DYNC1H1* locus using CRISPR/Cas9-mediated homology-directed repair. FW: Forward primer, RE: reverse primer. Right: PCR-based genotyping of the DHC-mClover KI cells. A single band of ∼5.1 kb confirmed homozygous insertion of the mClover cassette in clone #12. **c)** Quantification of the levels of the indicated CAAX constructs at the membrane during interphase and mitosis, showing that each construct was expressed and associated with the membrane continuously throughout the cell cycle. n = 30 cells ± SEM from three independent experiments. **d)** Live cell images showing endogenous dynein (DHC-mClover) recruitment by different levels of NuMA-N-mCherry-3xFlag-CAAX, indicating that the mitotic arrest phenotype was induced even in the presence of lower levels of NuMA-N. **e)** Representative live cell images of cortical dynein recruitment by the indicated adaptor constructs during prometaphase in the presence of 10 μM STLC. **f)** Quantification of membrane-associated DHC-mClover relative to its cytoplasmic signal. n = 30 cells ± SEM from three independent experiments; Brown-Forsythe and Welch ANOVA with Dunnett’s multiple comparisons tests. Scale bars represent 10 μm.

**Figure S2:**
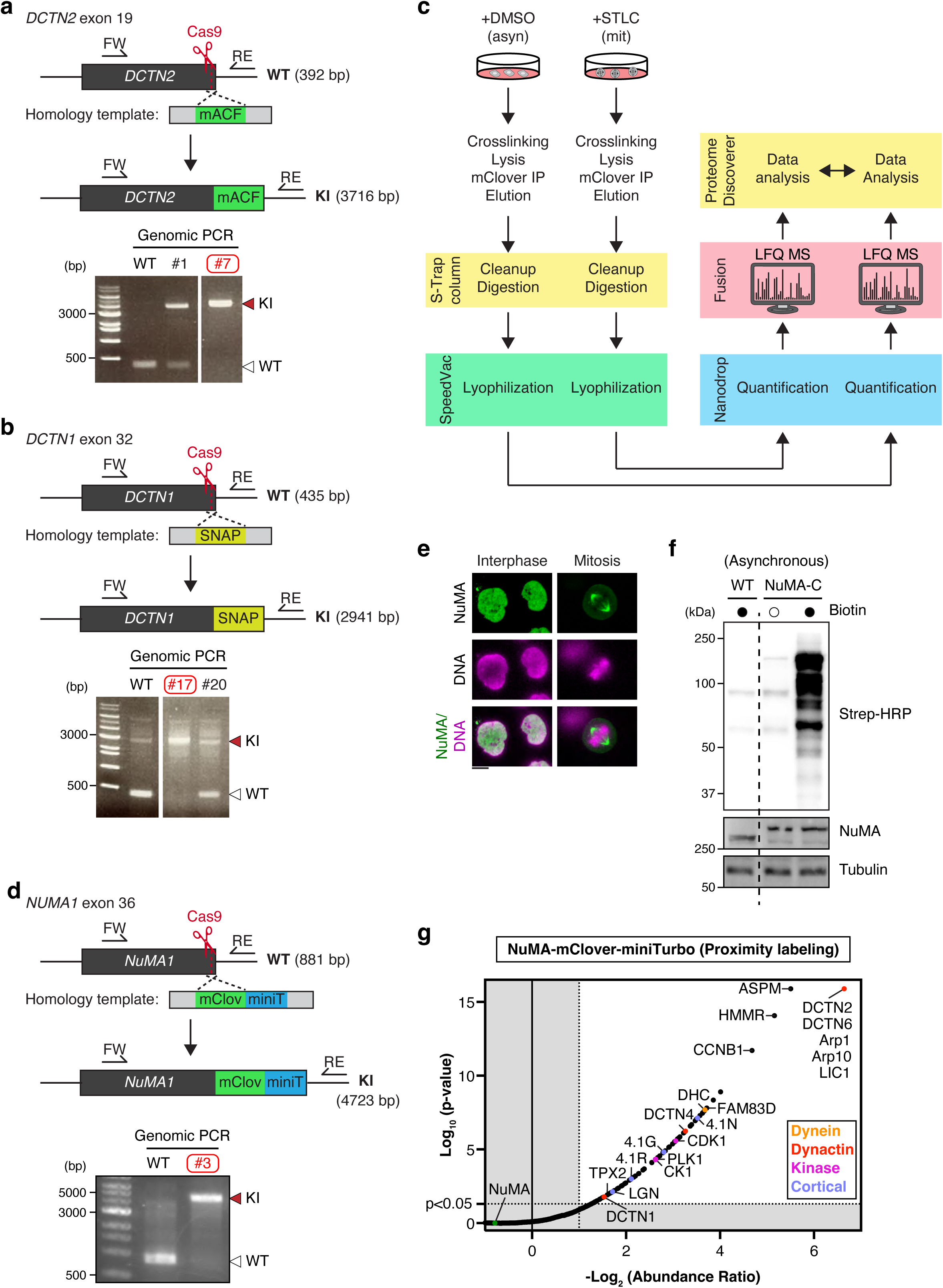
Proteomics approaches to identify mitosis-specific DDN interactors and modifications. **a)** Upper: Schematic depiction of the generation of DCTN2-mAID-mClover-3xFlag (DCTN2-mACF) HCT116 KI cells. The mACF cassette, together with a Neomycin selection marker (depicted as mACF for simplicity), were inserted at the C-terminus of the *DCTN2* locus using CRISPR/Cas9-mediated genome editing. FW: Forward primer, RE: reverse primer. Right: PCR-based genotyping of the DCTN2-mACF KI cells. A band of ∼3.7 kb and absence of WT ∼0.4 kb band confirmed homozygous insertion of the mACF cassette in clone #7. **b)** Upper: Schematic showing the generation of DCTN1-SNAP HCT116 KI cells (in DCTN2-mACF background) via CRISPR/Cas9-mediated genome editing. The SNAP cassette and a Blasticidine S selection marker were inserted at the C-terminus of the *DCTN1* locus. FW: Forward primer, RE: reverse primer. Right: PCR-based genotyping of the DCTN2-mACF KI cells. A band of ∼2.9 kb and absence of WT ∼0.4 kb band confirmed homozygous insertion of the SNAP cassette in clone #17. **c)** Diagram summarizing the proteomics and label-free quantification approach and procedure employed. IP: Immunoprecipitation, MS: Mass spectrometry. **d)** Upper: Schematic illustrating the creation of NuMA-mClover-miniTurbo HCT116 KI cells. The mClover-miniTurbo cassette and a Neomycin selection marker were inserted at the C-terminus of the *NUMA1* locus using CRISPR/Cas9-mediated homology-directed repair. FW: Forward primer, RE: reverse primer. Right: PCR-based genotyping of the NuMA-mClover-miniTurbo KI cells. A band of ∼4.7 kb confirmed homozygous insertion of the mClover cassette in clone #3. **e)** Live cell images showing nuclear and spindle accumulation of NuMA-mClover-miniTurbo during interphase and mitosis, respectively. **f)** Biotinylation assay showing the specificity and activity of NuMA-mClover-miniTurbo after addition of 50 μM biotin for 30 min. Cells were lysed with RIPA buffer and lysates were analyzed by immunoblotting. Strep-HRP was used to detect biotinylated proteins. **g)** Volcano plot showing mitosis-specific NuMA interactors identified by performing proximity labeling in asynchronously growing and prometaphase-arrested (STLC-treated) HCT116 NuMA-mClover-miniTurbo KI cells. Eluates from Streptavidin-agarose beads were analyzed by mass spectrometry followed by label-free quantification using the CHIMERYS search algorithm.

**Figure S3:**
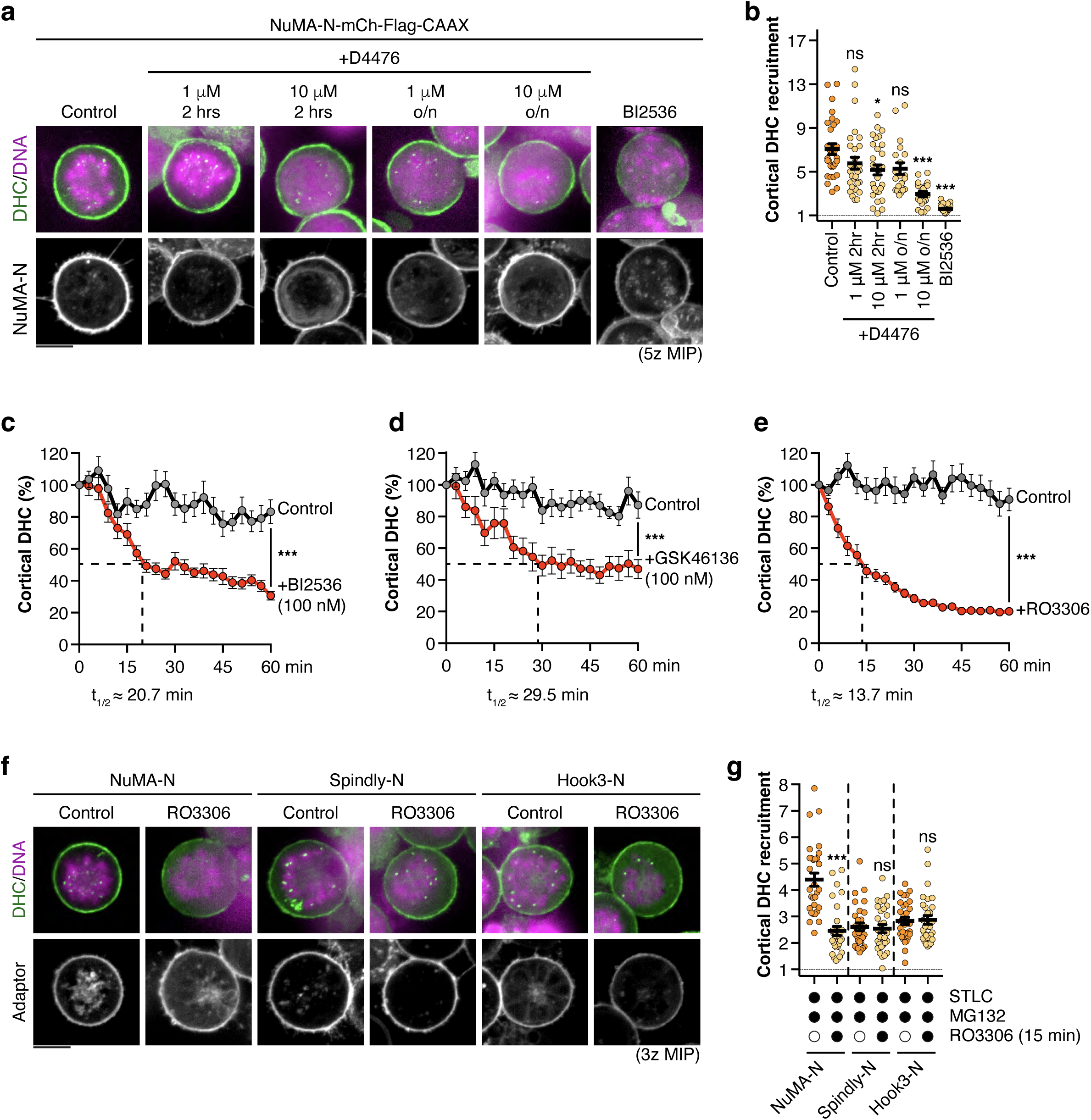
Analyzing effects of kinase inhibitors in the membrane-targeting assays. **a)** Representative live cell images of NuMA-N-mCherry-CAAX-dependent cortical dynein recruitment in mitotic Flp-In T-REx 293 DHC-mClover KI cells treated with the indicated doses of the CK1 inhibitor D4476 (2 h or overnight), or the PLK1 inhibitor BI2536 for 2 h. **b)** Quantification of relative DHC-mClover accumulation at the membrane in cells expressing membrane-tethered NuMA-N after incubation with the indicated concentrations of the CK1 inhibitor D4476 for 2 or ∼16 h (overnight). n = 30 cells ± SEM from three independent experiments; Brown-Forsythe and Welch ANOVA with Dunnett’s multiple comparisons tests. **c-e)** Time-lapse quantification of NuMA-dependent cortical DHC-mClover accumulation in Flp-In T-REx 293 cell after addition of either **c)** 100 nM BI2536, **d)** 100 nM GSK461364 or **e)** 9 μM RO3306. 0.1% DMSO was used as a control. Data were normalized to 100% at t = 0 min. n > 12 cells ± SEM from at least two independent experiments; unpaired Welch’s t-test on the final values (t = 60 min). **f)** Representative photographs of mitotic Flp-In T-REx 293 DHC-mClover KI cells expressing the indicated N-terminal adaptor constructs, treated with either 9 μM RO3306 or 0.1% DMSO for 15 min after pre-incubation with 20 μM MG132 for 1 hr. **g)** Quantification of cortical dynein. n = 30 cells ± SEM from two independent experiments; Brown-Forsythe and Welch ANOVA with Dunnett’s multiple comparisons tests. Scale bars represent 10 μm.

**Figure S4:**
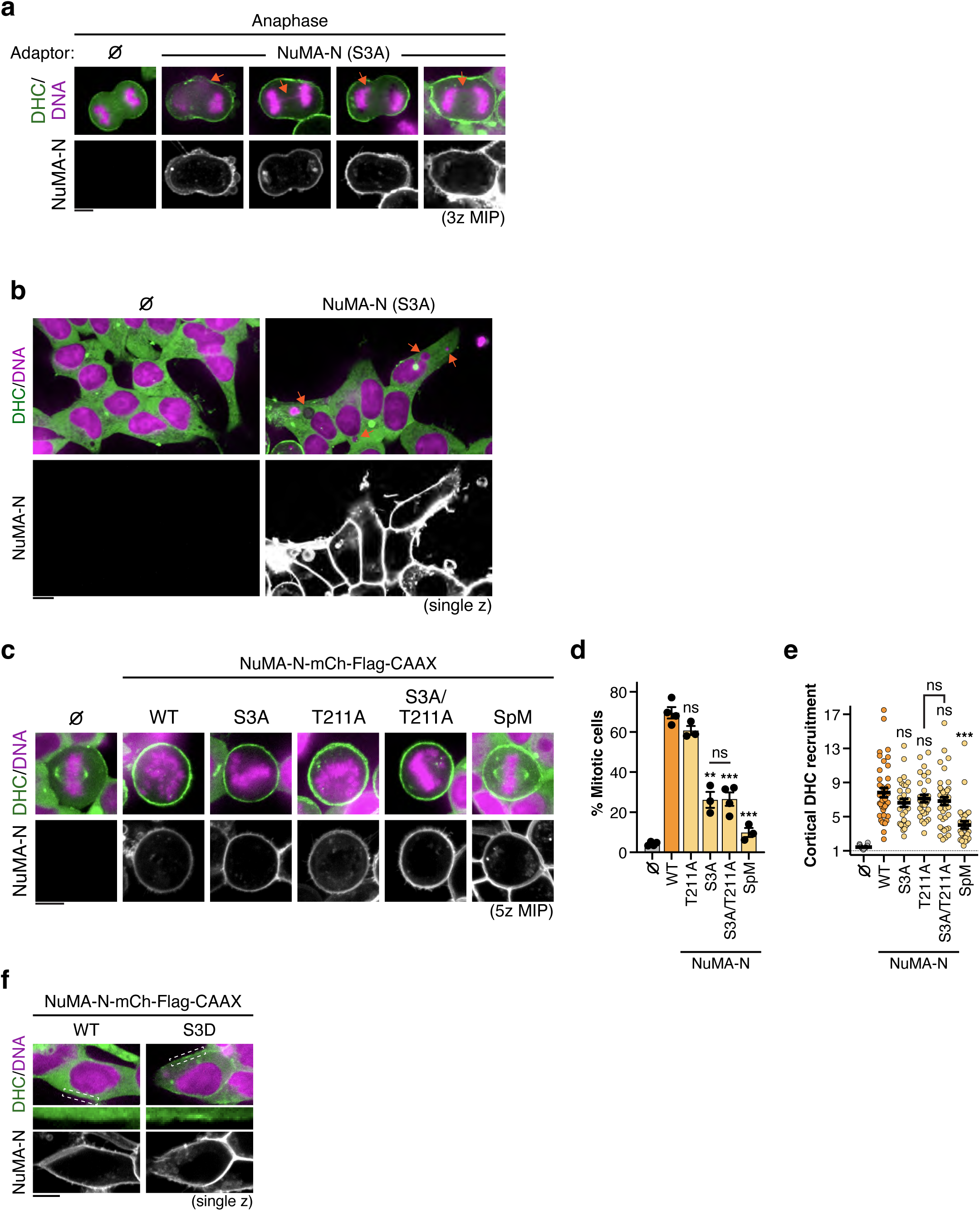
Mitotic and interphase phenotypes caused by membrane tethering of mutated NuMA-N fragments. **a)** Examples of anaphase entry as observed in Flp-In T-REx 293 DHC-mClover KI cells expressing membrane-tethered NuMA-N WT or S3A. Orange arrows highlight chromosome missegregation phenotypes. **b)** Live cell images of interphase Flp-In T-REx 293 DHC-mClover KI expressing NuMA-N S3A, showing the occurrence of micronuclei, pointed out by orange arrows. As cells expressing NuMA-N WT do not enter anaphase, we were unable to compare these with cells expressing NuMA-N S3A. **c)** Representative images showing DHC-mClover recruitment to the indicated membrane-tethered NuMA-N phosphomutants. **d)** Percentage of mitotic cells after expression of the displayed N-terminal NuMA mutants. **e)** Quantification of cortical dynein accumulation in cells expressing the indicated NuMA-N constructs. n = 30 cells ± SEM from three independent experiments; Brown-Forsythe and Welch ANOVA with Dunnett’s multiple comparisons tests. **f)** Live cell stills of interphase Flp-In T-REx 293 DHC-mClover KI expressing membrane-associated NuMA-N WT or S3D constructs, highlighting weak dynein signals at the membrane of selected cells expressing NuMA-N S3D. Scale bars represent 10 μm.

**Figure S5:**
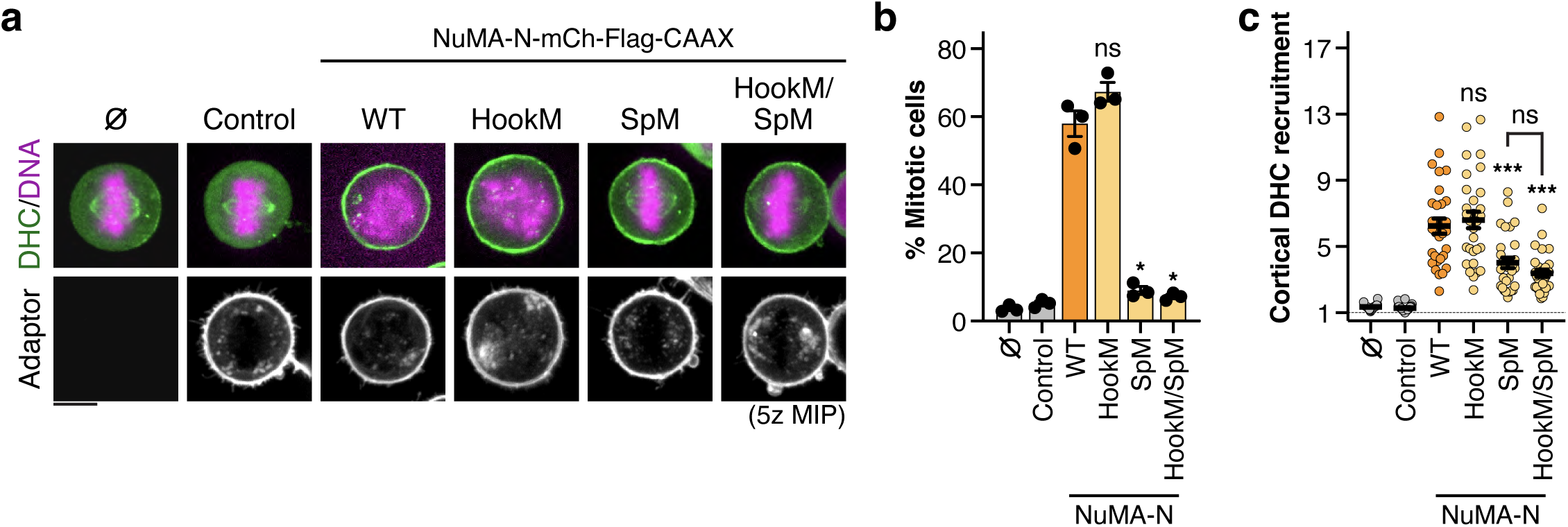
Mitotic phenotypes caused by membrane-tethering of N-terminal NuMA motif mutants. **a)** Representative live cell photographs of DHC-mClover recruitment to the indicated membrane-bound NuMA-N mutants tagged with mCherry-3xFlag-CAAX or mCherry-3xFlag-CAAX alone (Control). **b)** Quantification of the percentage of mitotic cells after expression of the displayed N-terminal NuMA mutants. **c)** Quantification of relative cortical DHC-mClover accumulation after expression of the displayed N-terminal NuMA mutants. n = 30 cells ± SEM from three independent experiments; Brown-Forsythe and Welch ANOVA with Dunnett’s multiple comparisons tests.

**Figure S6:**
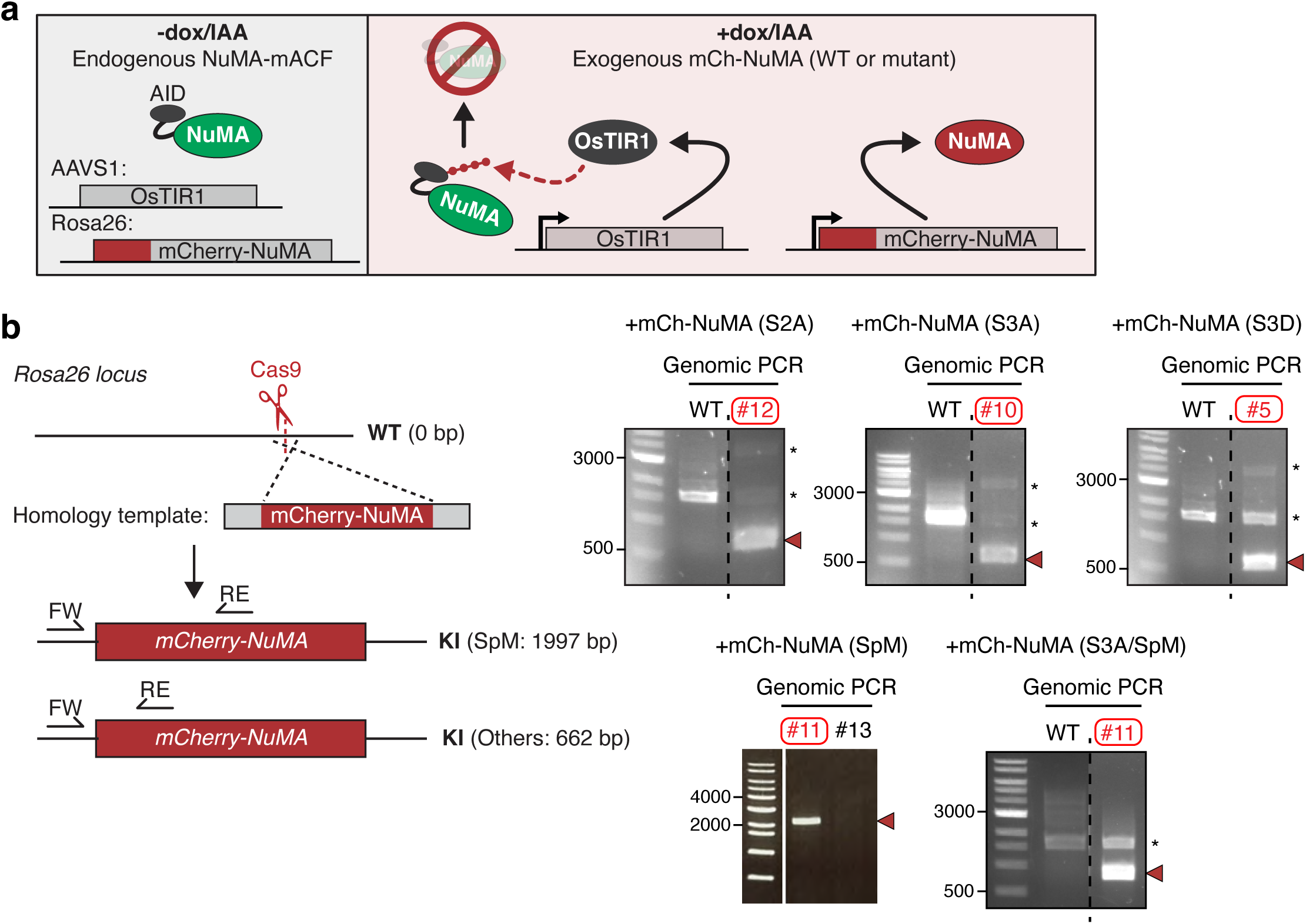
Cell line establishment for AID-mediated replacement of endogenous NuMA with NuMA mutants. **a)** Schematic representation of protein replacement strategy. Upon addition of Dox and IAA, endogenous NuMA-mACF is degraded while mCherry-NuMA constructs are expressed from the Rosa26 locus. **b)** Left: Diagram representing the generation of mCherry-NuMA rescue cell lines in HCT116 NuMA-mACF + DHC-SNAP double KI cells. The mCherry-NuMA cassettes (together with Hygromycin selection markers, depicted as mCherry-NuMA for simplicity), were inserted at the *Rosa26* safe harbor locus using CRISPR/Cas9-mediated homology-directed repair. FW: Forward primer, RE: reverse primer. Right: PCR-based genotyping of the mCherry-NuMA rescue cell lines. A band of ∼2 (SpM) or ∼0.7 (S2A, S3A, S3D and S3A/SpM) kb confirmed insertion of the mCherry-NuMA cassettes in the highlighted clones.

**Figure S7.**
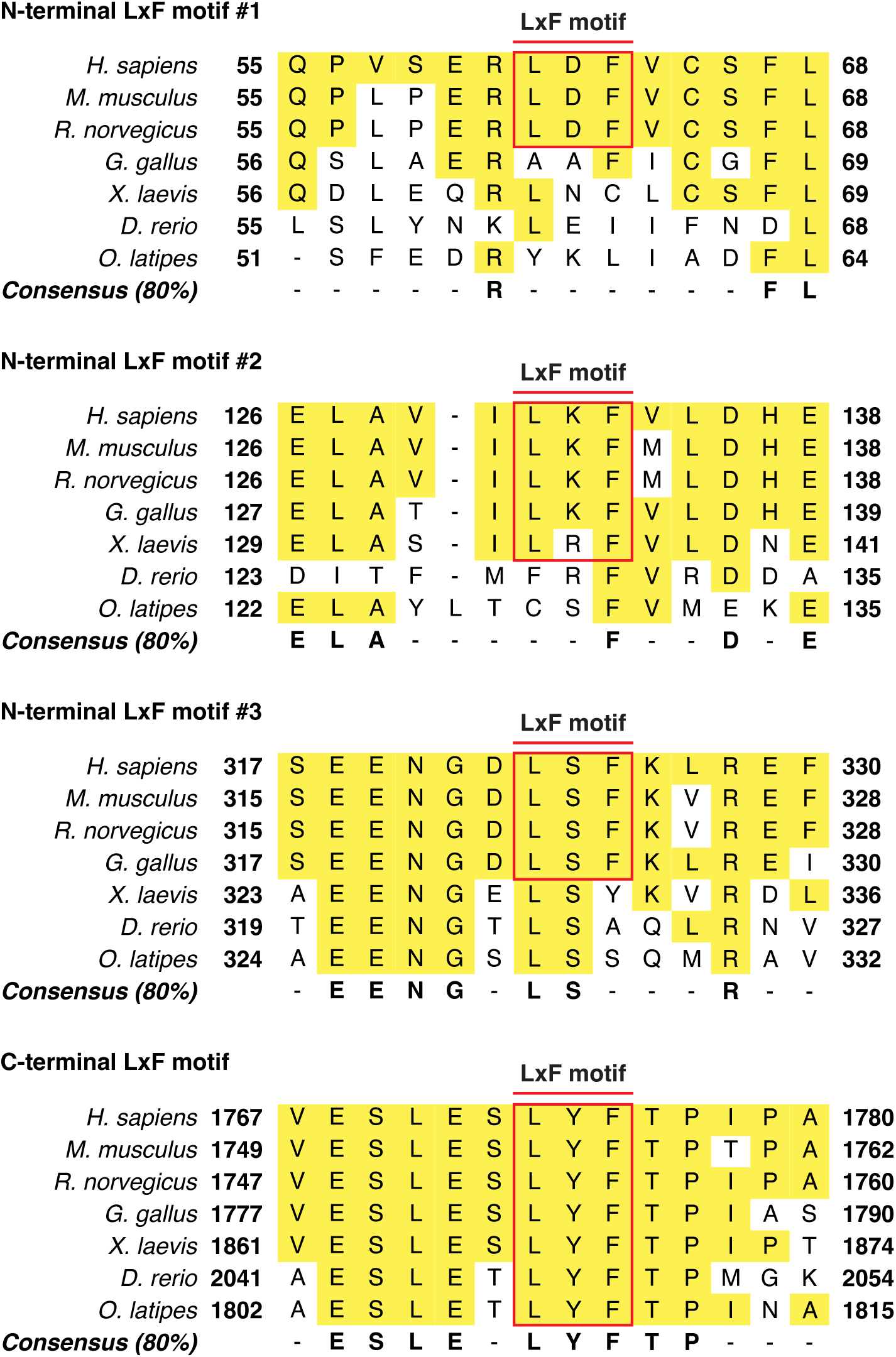
Sequence alignment of the Cyclin B-binding motifs on NuMA. Sequence alignment showing the conservation of LxF (Cyclin B-binding) motif embedded within the N- and C-termini of NuMA.

## References

1. Prosser, S. L. & Pelletier, L. Mitotic spindle assembly in animal cells: a fine balancing act. Nat Rev Mol Cell Biol 18, 187–201 (2017).

2. Valdez, V. A., Neahring, L., Petry, S. & Dumont, S. Mechanisms underlying spindle assembly and robustness. Nat Rev Mol Cell Biol 24, 523–542 (2023).

3. Petry, S. Mechanisms of Mitotic Spindle Assembly. Annu Rev Biochem 85, 659–683 (2016).

4. Santaguida, S. & Amon, A. Short- and long-term effects of chromosome mis-segregation and aneuploidy. Nat Rev Mol Cell Biol 16, 473–485 (2015).

5. Radulescu, A. E. & Cleveland, D. W. NuMA after 30 years: the matrix revisited. Trends Cell Biol 20, 214–222 (2010).

6. Kiyomitsu, T. & Boerner, S. The Nuclear Mitotic Apparatus (NuMA) Protein: A Key Player for Nuclear Formation, Spindle Assembly, and Spindle Positioning. Front Cell Dev Biol 9, 728 (2021).

7. Serra-Marques, A. et al. The mitotic protein NuMA plays a spindle-independent role in nuclear formation and mechanics. Journal of Cell Biology 219, e202004202 (2020).

8. Ray, S. et al. A mechanism for oxidative damage repair at gene regulatory elements. Nature 609, 1038–1047 (2022).

9. Chinen, T. et al. NuMA assemblies organize microtubule asters to establish spindle bipolarity in acentrosomal human cells. EMBO J 39, e102378 (2020).

10. Sun, M. et al. NuMA regulates mitotic spindle assembly, structural dynamics and function via phase separation. Nat Commun 12, 7157 (2021).

11. van Toorn, M., Gooch, A., Boerner, S. & Kiyomitsu, T. NuMA deficiency causes micronuclei via checkpoint-insensitive k-fiber minus-end detachment from mitotic spindle poles. Current Biology 33, 572–580.e2 (2023).

12. Hueschen, C. L., Kenny, S. J., Xu, K. & Dumont, S. NuMA recruits dynein activity to microtubule minus-ends at mitosis. Elife 6, e29328 (2017).

13. Okumura, M., Natsume, T., Kanemaki, M. T. & Kiyomitsu, T. Dynein–Dynactin–NuMA clusters generate cortical spindle-pulling forces as a multi-arm ensemble. Elife 7, e36559 (2018).

14. Gallini, S. et al. NuMA Phosphorylation by Aurora-A Orchestrates Spindle Orientation. Current Biology 26, 458–469 (2016).

15. Merdes, A., Heald, R., Samejima, K., Earnshaw, W. C. & Cleveland, D. W. Formation of Spindle Poles by Dynein/Dynactin-Dependent Transport of Numa. Journal of Cell Biology 149, 851–862 (2000).

16. Tsuchiya, K. et al. Ran-GTP Is Non-essential to Activate NuMA for Mitotic Spindle-Pole Focusing but Dynamically Polarizes HURP Near Chromosomes. Current Biology 31, 115–127.e3 (2021).

17. Silk, A. D., Holland, A. J. & Cleveland, D. W. Requirements for NuMA in maintenance and establishment of mammalian spindle poles. Journal of Cell Biology 184, 677–690 (2009).

18. Aslan, M. et al. Structural and functional insights into activation and regulation of the dynein-dynactin-NuMA complex. *bioRxiv* 2024.11.26.625568 (2024) doi:10.1101/2024.11.26.625568.

19. Colombo, S. et al. NuMA is a mitotic adaptor protein that activates dynein and connects it to microtubule minus ends. Journal of Cell Biology 224, e202408118 (2025).

20. Zhang, K. et al. Cryo-EM Reveals How Human Cytoplasmic Dynein Is Auto-inhibited and Activated. Cell 169, 1303–1314.e18 (2017).

21. Torisawa, T. et al. Autoinhibition and cooperative activation mechanisms of cytoplasmic dynein. Nat Cell Biol 16, 1118–1124 (2014).

22. Reck-Peterson, S. L., Redwine, W. B., Vale, R. D. & Carter, A. P. The cytoplasmic dynein transport machinery and its many cargoes. Nat Rev Mol Cell Biol 19, 382–398 (2018).

23. Roberts, A. J., Kon, T., Knight, P. J., Sutoh, K. & Burgess, S. A. Functions and mechanics of dynein motor proteins. Nat Rev Mol Cell Biol 14, 713–726 (2013).

24. Markus, S. M., Marzo, M. G. & McKenney, R. J. New insights into the mechanism of dynein motor regulation by lissencephaly-1. Elife 9, e59737 (2020).

25. Olenick, M. A. & Holzbaur, E. L. F. Dynein activators and adaptors at a glance. J Cell Sci 132, jcs227132 (2019).

26. Renna, C. et al. Organizational Principles of the NuMA-Dynein Interaction Interface and Implications for Mitotic Spindle Functions. Structure 28, 820–829.e6 (2020).

27. Terawaki, S., Yoshikane, A., Higuchi, Y. & Wakamatsu, K. Structural basis for cargo binding and autoinhibition of Bicaudal-D1 by a parallel coiled-coil with homotypic registry. Biochem Biophys Res Commun 460, 451–456 (2015).

28. Hoogenraad, C. C. et al. Mammalian Golgi-associated Bicaudal-D2 functions in the dynein– dynactin pathway by interacting with these complexes. EMBO J 20, 4041-4054–4054 (2001).

29. Schroeder, C. M. & Vale, R. D. Assembly and activation of dynein–dynactin by the cargo adaptor protein Hook3. Journal of Cell Biology 214, 309–318 (2016).

30. d’Amico, E. A., et al. Conformational transitions of the Spindly adaptor underlie its interaction with Dynein and Dynactin. Journal of Cell Biology 221, e202206131 (2022).

31. Urnavicius, L. et al. Cryo-EM shows how dynactin recruits two dyneins for faster movement. Nature 554, 202–206 (2018).

32. Horgan, C. P., Hanscom, S. R., Jolly, R. S., Futter, C. E. & McCaffrey, M. W. Rab11-FIP3 links the Rab11 GTPase and cytoplasmic dynein to mediate transport to the endosomal-recycling compartment. J Cell Sci 123, 181–191 (2010).

33. Gallisà-Suñé, N. et al. BICD2 phosphorylation regulates dynein function and centrosome separation in G2 and M. Nat Commun 14, 2434 (2023).

34. Kettenbach, A. N. et al. Quantitative Phosphoproteomics Identifies Substrates and Functional Modules of Aurora and Polo-Like Kinase Activities in Mitotic Cells. Sci Signal 4, rs5–rs5 (2011).

35. Compton, D. A. & Luo, C. Mutation of the predicted p34cdc2 phosphorylation sites in numa impair the assembly of the mitotic spindle and block mitosis. J Cell Sci 108, 621–633 (1995).

36. Kotak, S., Busso, C. & Gönczy, P. NuMA phosphorylation by CDK1 couples mitotic progression with cortical dynein function. EMBO J 32, 2517-2529–2529 (2013).

37. Sana, S., Keshri, R., Rajeevan, A., Kapoor, S. & Kotak, S. Plk1 regulates spindle orientation by phosphorylating NuMA in human cells. Life Sci Alliance 1, e201800223 (2018).

38. Polverino, F. et al. The Aurora-A/TPX2 Axis Directs Spindle Orientation in Adherent Human Cells by Regulating NuMA and Microtubule Stability. Current Biology 31, 658–667.e5 (2021).

39. Kotak, S., Afshar, K., Busso, C. & Gönczy, P. Aurora A kinase regulates proper spindle positioning in C. elegans and in human cells. J Cell Sci 129, 3015–3025 (2016).

40. Matsumura, S. et al. ABL1 regulates spindle orientation in adherent cells and mammalian skin. Nat Commun 3, 626 (2012).

41. Kotak, S., Busso, C. & Gönczy, P. Cortical dynein is critical for proper spindle positioning in human cells. Journal of Cell Biology 199, 97–110 (2012).

42. Chang, C.-C., Huang, T.-L., Shimamoto, Y., Tsai, S.-Y. & Hsia, K.-C. Regulation of mitotic spindle assembly factor NuMA by Importin-β. Journal of Cell Biology 216, 3453–3462 (2017).

43. Compton, D. A. & Cleveland, D. W. NuMA, a nuclear protein involved in mitosis and nuclear reformation. Curr Opin Cell Biol 6, 343–346 (1994).

44. Lydersen, B. K. & Pettijohn, D. E. Human-specific nuclear protein that associates with the polar region of the mitotic apparatus: Distribution in a human/hamster hybrid cell. Cell 22, 489–499 (1980).

45. Kiyomitsu, T., Hiroaki, M. & and Yanagida, M. Protein Interaction Domain Mapping of Human Kinetochore Protein Blinkin Reveals a Consensus Motif for Binding of Spindle Assembly Checkpoint Proteins Bub1 and BubR1. Mol Cell Biol 31, 998–1011 (2011).

46. Adam, S. A., Marr, R. S. & Gerace, L. Nuclear protein import in permeabilized mammalian cells requires soluble cytoplasmic factors. Journal of Cell Biology 111, 807–816 (1990).

47. van der Voet, M. et al. NuMA-related LIN-5, ASPM-1, calmodulin and dynein promote meiotic spindle rotation independently of cortical LIN-5/GPR/Gα. Nat Cell Biol 11, 269–277 (2009).

48. Du, Q. & Macara, I. G. Mammalian Pins Is a Conformational Switch that Links NuMA to Heterotrimeric G Proteins. Cell 119, 503–516 (2004).

49. Fulcher, L. J. et al. FAM83D directs protein kinase CK1α to the mitotic spindle for proper spindle positioning. EMBO Rep 20, e47495 (2019).

50. Branon, T. C. et al. Efficient proximity labeling in living cells and organisms with TurboID. Nat Biotechnol 36, 880–887 (2018).

51. Ma, L. et al. Requirement for Nudel and dynein for assembly of the lamin B spindle matrix. Nat Cell Biol 11, 247–256 (2009).

52. Vassilev, L. T. et al. Selective small-molecule inhibitor reveals critical mitotic functions of human CDK1. Proceedings of the National Academy of Sciences 103, 10660–10665 (2006).

53. Steegmaier, M. et al. BI 2536, a Potent and Selective Inhibitor of Polo-like Kinase 1, Inhibits Tumor Growth In Vivo. Current Biology 17, 316–322 (2007).

54. Görgün, G. et al. A novel Aurora-A kinase inhibitor MLN8237 induces cytotoxicity and cell-cycle arrest in multiple myeloma. Blood 115, 5202–5213 (2010).

55. Rena, G., Bain, J., Elliott, M. & Cohen, P. D4476, a cell-permeant inhibitor of CK1, suppresses the site-specific phosphorylation and nuclear exclusion of FOXO1a. EMBO Rep 5, 60-65–65 (2004).

56. Shuda, M. et al. CDK1 substitutes for mTOR kinase to activate mitotic cap-dependent protein translation. Proceedings of the National Academy of Sciences 112, 5875–5882 (2015).

57. Johnson, J. L. et al. An atlas of substrate specificities for the human serine/threonine kinome. Nature 613, 759–766 (2023).

58. Miller, M. L. et al. Linear Motif Atlas for Phosphorylation-Dependent Signaling. Sci Signal 1, ra2–ra2 (2008).

59. Shintomi, K., Masahara-Negishi, Y., Shima, M., Tane, S. & Hirano, T. Recombinant cyclin B-Cdk1-Suc1 capable of multi-site mitotic phosphorylation in vitro. PLoS One 19, e0299003-(2024).

60. Peters, J.-M. The Anaphase-Promoting Complex: Proteolysis in Mitosis and Beyond. Mol Cell 9, 931–943 (2002).

61. Krupina, K., Goginashvili, A. & Cleveland, D. W. Causes and consequences of micronuclei. Curr Opin Cell Biol 70, 91–99 (2021).

62. Kiyomitsu, T. & Boerner, S. The Nuclear Mitotic Apparatus (NuMA) Protein: A Key Player for Nuclear Formation, Spindle Assembly, and Spindle Positioning. Front Cell Dev Biol 9, (2021).

63. Natsume, T., Kiyomitsu, T., Saga, Y. & Kanemaki, M. T. Rapid Protein Depletion in Human Cells by Auxin-Inducible Degron Tagging with Short Homology Donors. Cell Rep 15, 210–218 (2016).

64. Raaijmakers, J. A., Tanenbaum, M. E. & Medema, R. H. Systematic dissection of dynein regulators in mitosis. Journal of Cell Biology 201, 201–215 (2013).

65. Firestone, A. J. et al. Small-molecule inhibitors of the AAA+ ATPase motor cytoplasmic dynein. Nature 484, 125–129 (2012).

66. Örd, M., Venta, R., Möll, K., Valk, E. & Loog, M. Cyclin-Specific Docking Mechanisms Reveal the Complexity of M-CDK Function in the Cell Cycle. Mol Cell 75, 76–89.e3 (2019).

67. Portegijs, V. et al. Multisite Phosphorylation of NuMA-Related LIN-5 Controls Mitotic Spindle Positioning in C. elegans. PLoS Genet 12, e1006291-(2016).

68. Zheng, Z., Wan, Q., Meixiong, G. & Du, Q. Cell cycle–regulated membrane binding of NuMA contributes to efficient anaphase chromosome separation. Mol Biol Cell 25, 606–619 (2013).

69. Kiyomitsu, T. & Cheeseman, I. M. Chromosome- and spindle-pole-derived signals generate an intrinsic code for spindle position and orientation. Nat Cell Biol 14, 311–317 (2012).

70. Gobran, M. et al. PLK1 inhibition delays mitotic entry revealing changes to the phosphoproteome of mammalian cells early in division. EMBO J 44, 1891-1920–1920 (2025).

71. Crncec, A. & Hochegger, H. Triggering mitosis. FEBS Lett 593, 2868–2888 (2019).

72. Barisic, M. et al. Spindly/CCDC99 Is Required for Efficient Chromosome Congression and Mitotic Checkpoint Regulation. Mol Biol Cell 21, 1968–1981 (2010).

73. Chaaban, S. & Carter, A. P. Structure of dynein–dynactin on microtubules shows tandem adaptor binding. Nature 610, 212–216 (2022).

74. Cho, N. H., Aslan, M., Yildiz, A. & Dumont, S. NuMA mechanically reinforces the spindle independently of its partner dynein. *bioRxiv* 2024.11.29.622360 (2024) doi:10.1101/2024.11.29.622360.

75. Cundell, M. J. et al. A PP2A-B55 recognition signal controls substrate dephosphorylation kinetics during mitotic exit. Journal of Cell Biology 214, 539–554 (2016).

76. Mochida, S., Maslen, S. L., Skehel, M. & Hunt, T. Greatwall Phosphorylates an Inhibitor of Protein Phosphatase 2Α That Is Essential for Mitosis. Science (1979) 330, 1670–1673 (2010).

77. Kiyomitsu, T. & Cheeseman, I. M. Cortical Dynein and Asymmetric Membrane Elongation Coordinately Position the Spindle in Anaphase. Cell 154, 391–402 (2013).

78. Romé, P. et al. Aurora A contributes to p150glued phosphorylation and function during mitosis. Journal of Cell Biology 189, 651–659 (2010).

79. Kumari, A. et al. Phosphorylation and Pin1 binding to the LIC1 subunit selectively regulate mitotic dynein functions. Journal of Cell Biology 220, e202005184 (2021).

80. Whyte, J. et al. Phosphorylation regulates targeting of cytoplasmic dynein to kinetochores during mitosis. Journal of Cell Biology 183, 819–834 (2008).

81. Sharma, A., Dagar, S. & Mylavarapu, S. V. S. Transgelin-2 and phosphoregulation of the LIC2 subunit of dynein govern mitotic spindle orientation. J Cell Sci 133, jcs239673 (2020).

82. Malumbres, M. & Barbacid, M. Cell cycle, CDKs and cancer: a changing paradigm. Nat Rev Cancer 9, 153–166 (2009).

83. van Toorn, M. et al. Active DNA damage eviction by HLTF stimulates nucleotide excision repair. Mol Cell 82, (2022).

84. Frankenfield, A. M., Ni, J., Ahmed, M. & Hao, L. Protein Contaminants Matter: Building Universal Protein Contaminant Libraries for DDA and DIA Proteomics. J Proteome Res 21, 2104–2113 (2022).

